# Combining population genomics with ancient DNA to understand island colonization history of the Madagascar turtle dove

**DOI:** 10.1101/2025.06.05.658009

**Authors:** Nisha Dwivedi, Sean P. Heighton, Romuald Laso-Jadart, Alexander J. F. Verry, Alba Nieto-Heredia, Pierre Lesturgie, Danielle Khost, Mylène Bohec, Timothy B. Sackton, Lounes Chikhi, Ludovic Orlando, Julian P. Hume, Guillaume Achaz, Catherine Thèves, Stefano Mona, Ben H. Warren

## Abstract

The Mascarene archipelago (Mauritius, Reunion and Rodrigues), characterized by first human arrival being recent, offers a unique setting to study species colonization. Here we use a combination of modern and ancient DNA data as a case study to investigate the recent colonization history of a species of concern in relation to conservation programs – the Madagascar turtle dove (*Nesoenas picturata*) on Mauritius and Reunion. We generated a reference genome and re-sequenced genomes from contemporary *N. picturata* populations, as well as genome-wide data from relevant subfossils. A combination of model-free inferences, site frequency spectrum (SFS) based demographic modelling, and analyses of population structure including that of subfossils indicate that *N. picturata* colonized both islands independently and naturally from Madagascar, long before human arrival. Summary statistics and SFS-based modeling reveal large effective population sizes (*Ne*) and high genetic diversity in island populations, conflicting with historical accounts of human-induced demographic collapse. Based on goodness-of-fit, genetic structure and diversity indices do not discriminate between two solutions, one of which posits large recent *Ne* and negligible translocation rates, while the other supports recent severe bottlenecks followed by high post-human translocation from Madagascar. Nonetheless, linkage disequilibrium provides stronger evidence for the latter scenario, which may also explain high genetic diversity. Both modern and ancient DNA data sources independently support the classification of *N. picturata* as native to both islands. Our findings highlight the importance of validating demographic models with multiple summary statistics, and potential of using a combination of different data sources to resolve colonization history in recent time.

## Introduction

Over the past century, globalization has driven an exponential increase in human-mediated species introductions (Bertelsmeier et al., 2025; Roy et al., 2024). Non-native species, which are introduced to new regions by humans either accidentally or intentionally, often pose significant threats to native biodiversity through predation, competition, and habitat destruction (Freed & Cann, 2009; Mooney & Cleland, 2001). As a result, their presence is frequently opposed by scientists and conservation managers (Simberloff et al., 2013). While preventing the introduction of non-native species remains the most effective strategy, eradication is commonly employed to mitigate adverse effects once established (Roy et al., 2024; Russell et al., 2017). Given the severity of the measures required to eliminate non-native species, determining a species’ biogeographic origin is crucial for appropriate management of natural environments (Buckley & Catford, 2016; Simberloff et al., 2012, 2013).

Biogeographic origins at an intraspecific scale can be inferred from genome-wide data using a variety of approaches. Multiple population genetic methods are available to test specific demographic scenarios, usually applied to timescales of 10^4^ years and more (Marchi et al., 2021). These methods summarize genetic data with different approaches that include the site frequency spectrum (SFS), linkage disequilibrium (LD), identity by descent and haplotype structure (Beichman et al., 2017; Marchi et al., 2021). Demographic parameters are inferred from these genetic summaries through likelihood maximization or Bayesian inference, such as approximate Bayesian computation (Beaumont, 2010; Beerli, 2006; Estoup et al., 2004; Excoffier et al., 2013). However, parameter estimates can be inaccurate, in particular when focusing on recent generations, which are often the target when investigating the spread of non-native species. Assessing goodness-of-fit is an important approach in ensuring the biological and temporal relevance of demographic models and involves extensive post-hoc simulations along with integration of diverse genomic information (Johri et al., 2021). Ancient DNA potentially provides a complementary approach to check inferences on recent timescales, especially concerning the colonization of new environments (Leonardi et al., 2017). The distinct characteristics of oceanic island archipelagos, such as their geographic isolation, relatively small sizes and young ages, facilitate inferences of lineage history (Losos & Ricklefs, 2009; Martin et al., 2021; Warren et al., 2015). These distinct characteristics, in combination with detailed historical, ecological and paleontological records, provide ideal contexts for evaluating the biological relevance of demographic models, making island archipelagos natural laboratories for developing integrated genetic approaches across varying timescales.

Notwithstanding the advantages afforded by the increasing availability of whole genomes, resolving demographic history on recent timescales, including the role of human interventions, remains a challenge for population genomics in most situations (Santiago et al., 2020). In this context, the Mascarene archipelago, part of the Madagascar biodiversity hotspot (Myers et al., 2000; Schlüter, 2016; Thébaud et al., 2009), provides a particularly valuable arena for distinguishing natural versus anthropogenic events in a species’ biogeographic history. Unlike other biodiverse archipelagos worldwide, it was uninhabited prior to well-documented first human settlement approximately 400 years before present (YBP). Despite the archipelago being of volcanic origin (composed of three islands, Mauritius, Reunion, and Rodrigues) it has rich subfossil deposits (Hume, 2005), spanning the period before and after first human settlement. Such deposits document the severe pressure experienced by the native fauna since first human arrival, demonstrating that 50-60% of the native vertebrate fauna has since become extinct, including the dodo (Cheke & Hume, 2008; Probst & Brial, 2002). Extinctions in the Mascarenes are partly driven by the numerous introductions of non-native species (Cheke, 1987; Cheke & Hume, 2008). The negative effects of introduced species are even more potent on islands than on continents, especially those recently colonized by humans (Russell et al., 2017). Mascarene birds are particularly affected, with 17 endemic species being threatened (Hume, 2013). Another important issue for conserving the remaining Mascarene avifauna is that, while the propensity for hybridization varies across vertebrates, it is high in birds (Grant & Grant, 1997; Warren et al., 2012). For example, in the neighbouring Seychelles islands, hybridization between the native Seychelles turtle dove (*Nesoenas picturata rostrata*) and introduced *N. picturata* has resulted in the native species having the same phenotype as the invader (Rhymer & Simberloff, 1996).

*N. picturata* is unanimously considered native to Madagascar, and closely related to the endangered Mauritian pink pigeon (*N. mayeri*) (Cheke, 2005; Hume, 2011; Johnson et al., 2001). Subfossil evidence from Mauritius and Reunion suggests that native populations, corresponding to or affiliated with *N. picturata* once existed on these islands. Some of these subfossils, with more robust pelvic elements, have been used to propose the existence of an endemic but extinct Mauritian turtle dove – *N. cicur* (Hume, 2011). However, the colonization history of *N. picturata* across the archipelago remains unclear. Authorities disagree on whether the living populations are native to Mauritius and Reunion, or if the species was reintroduced following human settlement and the extinction of the original native populations. Despite uncertainty in the colonization history of *N. picturata*, management differs between Mauritius and Reunion. In Mauritius, the species is considered invasive and culled in areas where it overlaps with the pink pigeon. In Reunion, the species has been protected since 1983 after a population bottleneck in the 1990s (Barré, 1983; Probst, 2002). Currently, conservationists in Reunion face pressure from hunting lobbies to reconsider its protected status.

We generated whole genomes of modern individuals from Madagascar, Mauritius, and Reunion, as well as genome-wide data of subfossils from Mauritius. We investigated population structure, estimated genetic diversity and runs of homozygosity (ROH), and propose explicit demographic scenarios inferred using a coalescent framework that fits the observed SFS. We check the validity of our most likely scenario by modeling a posteriori the expected pattern of linkage disequilibrium, which was not used in the inferential procedure.

Our primary aim is to determine whether *N. picturata* is native or introduced in Mauritius and Reunion by estimating the route and timing of colonization of the Mascarenes. Second, we investigate the pre-versus post-human contribution of migration between landmasses to the modern *N. picturata* populations. Third, we test whether humans have had a positive or negative impact on the genetic diversity of the Mascarene island populations. Understanding the role of past human intervention in the distribution of a species is a globally-recurrent problem in conservation biology (Gallardo et al., 2015). We present a case study in resolving human contribution to colonization history, making a comparison between responses on recent timescales from ancient DNA sources with those from an integration of historical records, diverse modern genomic statistics and population genetics modelling. As such, our results have importance beyond the immediate management strategies of Mascarene conservation organizations. We demonstrate the potential of integrating multiple independent lines of molecular ecological evidence to uncover the recent colonization history of species of conservation concern (García-Verdugo et al., 2019; Theissinger et al., 2023).

## Methods

### Reference genome

We assembled a *de novo N. picturata* genome from an individual blood-sampled in 2022 at Plaine Lièvre, Black River Gorges National Park (Mauritius). Blood was flash-frozen in liquid nitrogen and high molecular weight (HMW) DNA was extracted using a Qiagen MagAttract HMW DNA kit (Qiagen, Hilden, Germany), extending incubation time and heating the elution buffer to improve DNA concentrations. See S1 for details of long-read PacBio sequencing and genome assembly.

### Modern sampling and genome resequencing

Our sampling covers the breadth of the species’ range on Mauritius (N=9) and Reunion (N=10), as well as geographically disparate regions of Madagascar (N=3), totaling 22 resequenced genomes (Figure 1). Twelve individuals were blood-sampled in the field in Mauritius (during 2020 and 2022) and Reunion (2022). Seven Reunion tissue samples come from the bird rehabilitation center of the Société d’Etudes Ornithologiques de La Réunion (SEOR). Three Madagascar samples were loaned by the Field Museum of Natural History. DNA was extracted using the Qiagen MagAttract HMW DNA kit (Qiagen, Hilden, Germany). Sequencing was performed by the IGenSeq core facility (Institut du Cerveau, Paris) on a NovaSeq 6000 sequencer.

**Figure 1.**
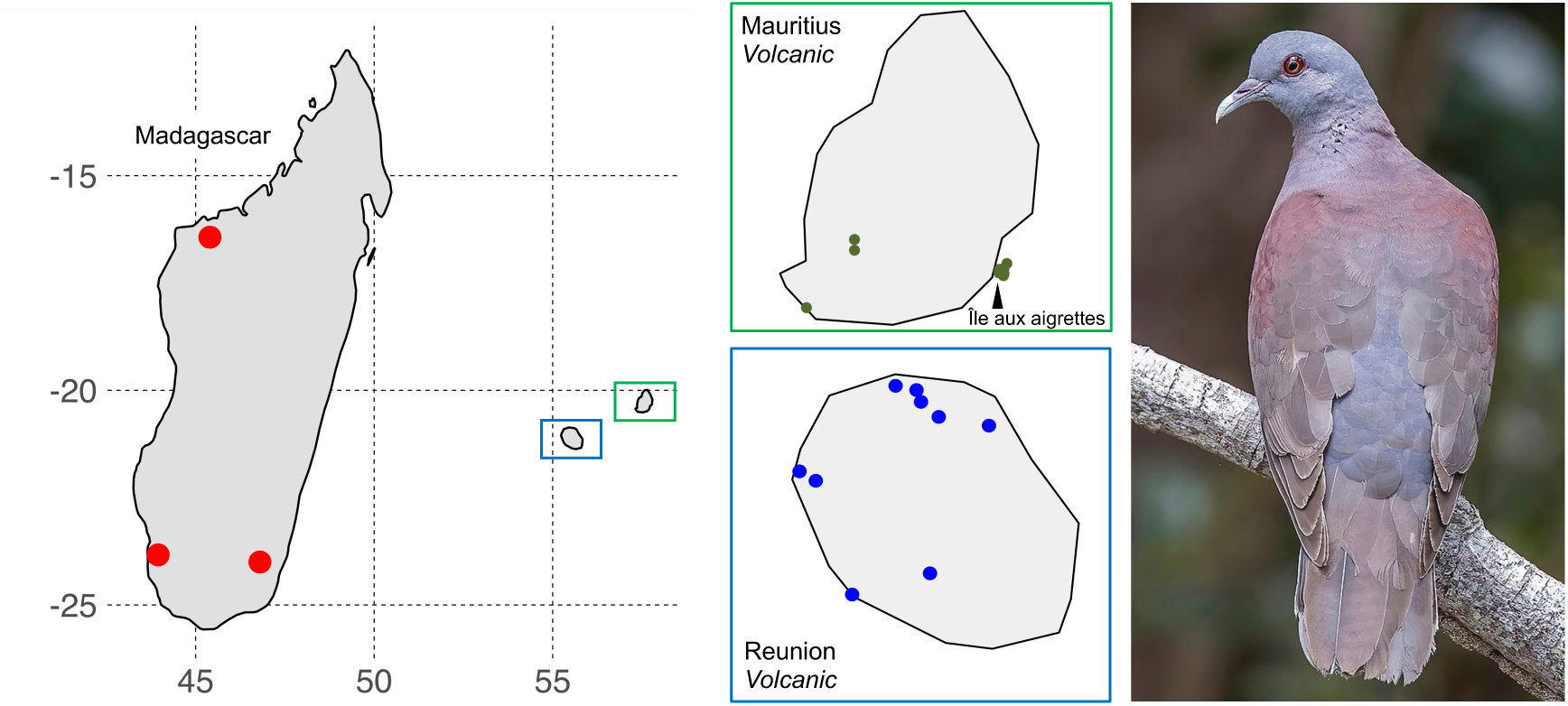
Sampling of *N. picturata* across the Mascarenes. Dots represent individual samples. Colours correspond to the population of origin: red= Madagascar; green= Mauritius; blue= Reunion. Italics indicates the geological nature of islands. Photo credits: CC Charles J. Sharp.

### Subfossil sampling and sequencing

Four *Nesoenas* subfossils from Mauritius morphologically identified as *N. picturata* or *N. cicur* were, with permission, destructively sampled for DNA (Table S1). Three were obtained from the collection of J.P. Hume and one was from the MNHN paleontology collection. One subfossil originates from an anoxic marsh (Mare aux Songes site) while the other three come from caves and boulder screes (Le Pouce and Le Morne sites; Table S1). All ancient DNA molecular work was carried out in the ancient DNA facilities of the Centre for Anthropobiology and Genomics of Toulouse, following strict experimental standards to avoid and monitor contamination, including use of disposable personal protection equipment, positive air pressure, bleach/UV surface and instrument decontamination, and negative control blanks. PCR amplification, purification, and quantification as well as DNA capture and preparation for sequencing were carried out in post-PCR laboratories that are located in a building physically isolated from the ancient DNA facilities. After gentle surface abrasion using a drill to remove potential contaminants, up to 100 mg of bone was powdered with the Mixel Mill 200 (Retsch) micro-dismembrator. DNA extraction using 30-50 mg of bone powder followed Gamba et al. (2016) and Fages et al. (2019). See S2 for details of ancient DNA methods, including shotgun library preparation, HyRAD probe development, hybridization capture and sequencing.

### Read mapping and variant calling of modern samples

Raw reads were processed with Trimmomatic v.0.39 to trim adaptors and remove low-quality segments, using default settings except for a minimum read length of 100 base pairs (bp) (Bolger et al., 2014). Remaining reads were mapped to the reference genome with bwa-mem v.0.7.17 (Li & Durbin, 2009). Alignments were sorted and indexed with SAMtools v.1.14 (Danecek et al., 2021). PCR duplicates were removed using Picard’s *MarkDuplicates* tool (https://broadinstitute.github.io/picard/). Variant calling was performed on the 32 scaffolds over 10M bp (83% of the genome) following GATK best practices, using GATK v4.2.6.1 (McKenna et al., 2010). BAM files were input into *HaplotypeCaller* using the *-ERC BP_RESOLUTION* flag (DePristo et al., 2011; Van Der Auwera et al., 2013). Resulting gVCF files were combined with *GenomicsDBImport* and jointly called using *GenotypeGVCF* with the *--include-non-variant-sites* flag to allow for the scaling of genetic diversity (DePristo et al., 2011; Van Der Auwera et al., 2013). We used VCFtools v0.1.16 to compute mean depth, quality, missingness, and inbreeding coefficient per site/individual to ensure data integrity (Danecek et al., 2021). Repetitive elements were removed from the VCF using bedtools v.2.30 *intersect* and the BED file described earlier (Quinlan & Hall, 2010).

VCF files were filtered differently for population structure and demographic analyses. For population structure, raw VCF files were filtered with VCFtools v.0.1.16 (Danecek et al., 2021) to keep only biallelic single-nucleotide polymorphisms (SNPs), removing SNPs with: a) genotypic depth <10 or >60; b) minor allele frequency (MAF) < 0.05; c) >20% missing genotypes; d) quality score <30. SNPs heterozygous in more than 80% of individuals were also removed using a custom R script to avoid repetitive region bias. For demographic analyses, which are sensitive to low-frequency variants, raw VCF files were filtered using GATK v.4.2.6.1’s *VariantFiltration* tool (DePristo et al., 2011; McKenna et al., 2010; Van Der Auwera et al., 2013), to remove potential sex chromosomes (scaffold 8 and 35), and discard sites with: a) quality score <30; b) genotypic depth <10 or >60; c) any missing genotype.

### Read mapping and variant calling of subfossil samples

To account for the highly fragmented nature of ancient DNA, the base modifications that accumulate post-mortem and the risk of contamination, we used a different bioinformatic pipeline for the subfossil samples. Raw reads were trimmed and filtered using AdapterRemoval v.2.3.3 (Schubert et al., 2016). Reads contained *Illumina* adaptors and seven bp internal barcodes at their extremities. *Illumina* adaptors were trimmed by setting a minimum adaptor overlap of three base pairs (bp) and a maximum mismatch fraction of 1/5. Internal barcodes were trimmed by setting the *-trim3p* and *-trim5p* flags to seven bp. We removed reads under 25 bp and stretches of low quality with ambiguous bases. Trimmed pair-end and single-end reads were processed separately to maximize the number of mapped reads. In both cases, reads were mapped to the reference genome using the bwa-aln algorithm with a minimum error rate of 0.1 and a seed length of 30 (Li & Durbin, 2009). Resulting SAM files were combined with bwa-sampe and bwa-samse (Li & Durbin, 2009). Additional filtering was applied to remove reads under 30 bp in length and 30 in mapping quality score. PCR duplicates were removed using the *markdup* tool in Samtools v.1.18 (Danecek et al., 2021). Pair-end and single-end BAM files were combined with Samtools v.1.18 *merge* (Danecek et al., 2021). DNA damage patterns were assessed with mapDamage v.2.2.2 and PMDtools v.0.60 (Jónsson et al., 2013; Skoglund et al., 2014). Due to minimal overlap between subfossil sequences, we computed the genotype likelihoods of each subfossil sample with the modern samples (previously obtained) separately. Genotype likelihoods were computed using the SAMtools (*-GL* 1) method implemented in ANGSD v.0.935 (Korneliussen et al., 2014). Only significant polymorphic sites (p-val<10-7) were retained. Tests showed that the patterns observed with a minimum depth of three were conserved at higher depths. Therefore, to ensure that we called sites with sufficient coverage in the subfossil samples, we set the *-minInd* flag to the total number of individuals and the *-setMinDepthInd* flag to three.

### Population structure

#### Modern samples

We investigated population structure using principal component analysis (PCA) using the *pcadapt* package v.4.3.3 in R (Privé et al., 2020). For each population pair (Mauritius-Reunion; Madagascar-Mauritius; Madagascar-Reunion), we estimated pairwise Fst with Reynold’s estimator (Reynolds et al., 1983), assessing significance through 1000 permutations in which individual sampling sites were randomized and calculating the number of re-sampled values larger that the observed value. We also used the sNMF clustering algorithm implemented in the *LEA* R package (Frichot & François, 2015). This program infers the number K of ancestral populations and estimates individual ancestry proportions. We ran 10 replicates for each value of K from 1 to 8 and investigate the cross-entropy curve to determine the most likely solution(s).

#### Subfossil samples

We used PCAngsd v.0.99 (Meisner & Albrechtsen, 2018) to perform PCA on subfossil and modern samples based on the previously calculated genotype likelihoods. Similarly, we computed distance matrices for each subfossil and the modern samples using the identity-by-state coefficient implemented in ANGSD (Meisner & Albrechtsen, 2018).

### Genetic diversity and ROH

Site frequency spectrum were computed from VCFs filtered for demography using a Python script (https://github.com/PierreLesturgie), from which we derived nucleotide diversity (θπ) (Tajima, 1983), Watterson’s theta (θw) (Watterson, 1975), and Tajima’s D (Tajima, 1989), normalizing θπ and θw by the total number of sites to obtain per site estimates of genetic diversity. These computations were performed over the entire genome and for individual scaffolds. We also calculated genome-wide individual observed heterozygosity (H_O_) from the sum of heterozygous positions in each individual scaled by the total number of sites.

ROH were estimated using the *--homozyg function* in PLINK v.1.90. This function uses sliding windows to identify consecutive homozygous segments. The input was a VCF filtered as for structure, but without the MAF filter. We did not apply LD or MAF filtering as it is discommended in the case of high quality datasets (Foote et al., 2021; Martin et al., 2023; Meyermans et al., 2020). Following Meyermans et al. (2020), we set parameters according to SNP density (0.2 kb/SNP) and data quality, maintaining consistent thresholds across all populations. The sliding window size was set to 300kb to detect both ancient and recent ROH. The thresholds used were: a) Minimum 50 SNPs per window (*--homozyg-snp);* b) Minimum SNP density of one per 50 kb (*--homozyg-density*); c) Minimum scanning window size of 50 SNPs (*--homozyg-window-snp*); d) Maximum gap of 200 kb between SNPs (--*homozyg-gap*); e) Maximum 2% heterozygous SNPs per segment (--*homozyg-window-het*); f) Maximum five missing genotypes per segment (--*homozyg-window-missing*).

### Demographic inference

We reconstructed the variation in the coalescent rate over time by comparing two methods known to perform differently as evolutionary time elapsed increases (Spence et al., 2018; Terhorst et al., 2017).

When panmixia cannot be assumed, the pairwise sequentially Markovian coalescent (PSMC v.0.6.5; Li & Durbin, 2011) remains a useful tool, harboring information on population structure or divergence (Lesturgie et al., 2022; Mazet et al., 2016). We created a consensus sequence on the reference genome for each individual with BCFtools v.1.11 *mpileup* and its *-c* flag. Resulting BCF files were converted to FASTQ using the *vcfutils.pl* perl script. We kept reads with depths ranging from 10 to 50, and with a minimum consensus base quality of 30. FASTQ files were converted to FASTA with the *fq2psmcfa* command. A bin size of 50 (*-s* flag) was used to account for the large SNP density (default values are specific to humans). To run PSMC v.0.6.5 we set the *-t* flag to 40, the *-r* flag to four and the *-p* flag to “95*2+4+6”. We changed the default time vector (*-p*) after tests prompted by Mather et al., (2020) showed that it reduces a bias caused by a low number of recombination events in recent time intervals. We bootstrapped variations in *Ne* by splitting the genome into 500,000 bins with the *splitfa* tool, and ran PSMC 100 times on randomly sampled bins.

The SMC++ v.1.15.2 program also uses a sequential Markovian coalescent approach to estimate the distribution of coalescence times, but leverages the SFS for better accuracy in recent times (Spence et al., 2018; Terhorst et al., 2017). We converted the VCF filtered for demography to SMC++ format with the *vcf2smc* tool (https://github.com/popgenmethods/smcpp). We looped through individuals, setting each one as the “distinguished” individual. The *estimate* command was used to compute the composite likelihood and the coalescence rate variations over time setting the *--knots* flag to 40.

For all historical demographic analyses, we used a generation time of 3.7 years (Seal & Bruford, 1991) and a substitution rate of 5.2x10^-9^ per site per generation (Shapiro et al., 2013). Due to the lack of specific estimates for the *N. picturata,* and given that genomic properties are often conserved in birds (Kawakami et al., 2014; Singhal et al., 2015; Zhang et al., 2014), we used values derived from studies of the congeneric pink pigeon (*Nesoenas mayeri*) and domestic rock pigeon (*Columba livia*). The pink pigeon is the sister species of *N. picturata*, with which it shares similar life history traits (Johnson et al., 2001; Shapiro et al., 2013).

### Demographic modeling

We performed coalescent-based maximum likelihood (ML) analyses using *fastsimcoal2* v.2709 (Excoffier et al., 2013; Marchi et al., 2021) to infer the most likely scenario for the historical demography of *N. picturata*. This method utilizes the pairwise two-dimensional SFS (2D-SFS) to estimate the parameters of given demographic scenarios. We generated 2D-SFS for each pair of populations from a VCF filtered for demography. We tested eight models, progressively increasing complexity, and compared each to the previous most likely model.

First, we tested several population tree topologies and colonizations sequence using a series of six models (Figure S1). To narrow the scope of models for testing, we accounted for geographical factors and population structure analysis results. Specifically, due to the distances between islands and the observed genetic structure, Madagascar was set as the ancestral deme in all but one of the tree topology models. We considered six models: i) MOD1, the population of Madagascar results from admixture between the populations of Mauritius and Reunion, included to test whether the sNMF clustering could be explained by this scenario; ii) MOD2, independent colonizations from Madagascar, with Mauritius colonized before Reunion; iii) MOD3, independent colonizations from Madagascar, with Reunion colonized before Mauritius; iv) MOD4, dependent colonizations, with the population of Reunion originating from Mauritius; v) MOD5, dependent colonizations, with the population of Mauritius originating from Reunion; and vi) MOD6, independent colonizations from Madagascar, with no fixed timing for the colonization events. Second, we assessed gene flow by incorporating asymmetric migration into the most likely tree topology, creating the seventh model, MOD6_MIG. Third, we evaluated changes in the *Ne* of Mauritius and Reunion, along with recent translocations from Madagascar to both islands, leading to the final model, MOD6_MIG_TR. The introduction of these additional parameters were guided by demographic inferences and historical records that suggest changes in *Ne* and frequent translocations of *N. picturata* individuals from Madagascar to Mauritius and Reunion (Cheke & Hume, 2008; Probst, 2002).

Each model was independently run 100 times (200 times for the final, most likely model), each with 40 expectation conditional maximization (ECM) cycles and 100,000 coalescent simulations for SFS estimation. Likelihood was evaluated excluding classes of the 2D-SFS with less than 10 occurrences. The models and their likelihood estimation were executed using the *fastsimcoal2* command: -n 100000 -m -M -L 40 -C 10. The specified range for each parameter is provided in Table S3. Models were evaluated using the Akaike Information Criterion (AIC). We also ensured that the likelihood distributions from 100 simulations under ML parameters did not overlap with those of previous models. For the most likely model, we performed parametric bootstraps to calculate the 95% confidence intervals of ML parameters for two runs of interest. For each run, this process involved simulating the model 100 times to generate 2D-SFS, running 100 independent parameter estimations for each simulated 2D-SFS (as done for the empirical data), and using the resulting parameter distributions to compute the 5% and 95% quantiles.

### Goodness-of-fit of the most likely model

#### Simulated structure and diversity

To evaluate the goodness-of-fit for two runs of the most likely model, we simulated the longest scaffold of the genome using the ML parameters from the parametric bootstraps of each run to estimate uncertainty. We set a per generation recombination rate of 10^-8^. Genotype tables were simulated using the following command options in *fastsimcoal2*: -n 100 -I -G -g -s0 -x -k 10000000. Using a custom bash script, we converted the genotype tables into VCFs and : i) computed the folded SFS from which we obtained θπ, θw, and Tajima’s D; ii) filtered out SNPs with a MAF lower than 0.05 and explored population structure through PCA, sNMF clustering and pairwise Fst.

#### IICR and LD statistics

We simulated 10^7^ independent pairwise times to most recent common ancestor (TMRCA) values for a pair of haploid chromosomes (*T_2_*) running *fastsimcoal2* with the *recordMRCA* flag on given demographic parameters. We repeated this process using the ML parameters from each parametric bootstrap of the most likely model for both runs of interest. We established a vector of 600 logarithmically spaced time points between 20 and 4.5x10^5^ generations in the past to obtain the cumulative density function of *T_2_*, F(*T_2_*) using a custom python script. See S3 for how the IICR and LD decay is obtained.

We followed the authors’ recommendations to compare theoretical with observed LD decay. We computed LD statistics for each population within specified recombination distance bins across 100 distant regions of 1M bp. These regions were obtained from a VCF file filtered for demography. The mid-points of the genetic distance bins (cM) are: 5e^-7^, 1.5e^-6^, 3.5e^-6^, 7.5e^-6^, 1.5e^-5^, 3.5e^-5^, 7.5e^-5^,1.5e^-4^, 3.5e^-4^, 7.5e^-4^. We excluded Madagascar due to low sample size. LD statistics, specifically 𝔼[D^2^] and 𝔼[D_z_], were calculated for all SNP pairs within each recombination distance bin for each population using the *compute_ld_statistics* function. The *means_from_region_data* function was used to compute the average LD statistics across all regions. All statistics were normalized by 𝔼[𝜋_2_] of the Reunion deme following *moments.LD* guidelines (Ragsdale & Gravel, 2019). Confidence intervals were calculated for observed data using the *bootstrap_data* function, which generates 100 bootstrap replicates by randomly sampling the 1M bp regions with replacement. We calculated expectations of LD decay from the ML parameters of parametric bootstraps of each run, using the *LDstats* command.

## Results

### Data quality and filtering

#### Modern samples

The mean sequencing depth for the entire dataset is 23X, increasing to 36X for scaffolds longer than 10M bp. (Table S1). After filtering, 16,466,416 SNPs were retained for population structure analyses. Filtering for demography resulted in 10,897,013 to 12,739,591 SNPs per population.

#### Subfossil samples

Endogenous DNA content of samples ranged from 0.03-1.5% before capture, and 0.11-5.8% after capture (Table S1). The number of mapped reads per subfossil ranges from 31,024 to 2,719,895 (Table S2). The distribution of DNA strand lengths from MiniSeq shotgun sequencing are between 35 and 55, as expected for subfossils. Authenticity profiles from deep sequencing of the non-USER-treated libraries generated with mapDamage2 and PMD-tools do not indicate significant DNA damage, which is not untypical for subfossils. The overlap between subfossil sequences, excluding modern genomes, was only 1,030 bp at a depth of 3X. The number of variants shared between each subfossil and the modern dataset ranges from 2,811 to 680,209.

### Population structure

#### Modern samples

PCA shows three distinct clusters corresponding to individuals from the three sampled islands, Madagascar, Mauritius, and Reunion (Figure 2A). PC1, explaining 16.7% of the total variation in the dataset, unambiguously segregates the three islands, indicating a stronger differentiation within the Mascarenes (between the islands of Mauritius and Reunion) than between either Mascarene island and Madagascar. Variation within the Mauritius population is represented along PC1 and PC2. Hierarchical clustering based on p-distances yields similar results (Figure S2). Consistent with the PCA, the highest pairwise Fst is between Reunion and Mauritius (Fst = 0.15, p=0.001; Figure 2B). There is a slightly higher Fst between Madagascar and Mauritius (Fst=0.11, p=0.006) than between Madagascar and Reunion (Fst=0.09, p=0.004; Figure 2B).

**Figure 2.**
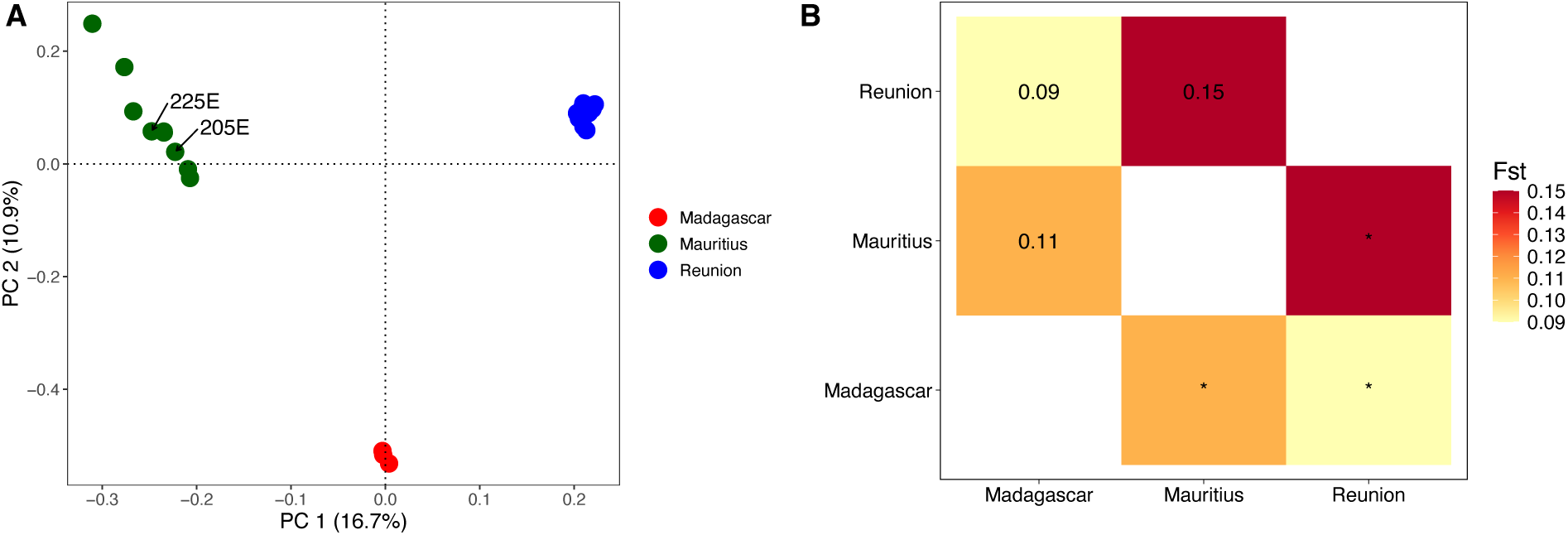
Population structure analyses with *N. picturata* genomes. **(A)** PCA. The axes represent the first two principal components (PC). Labels identify individual samples 205E and 225E. **(B)** Heatmap of the pairwise Reynold’s Fst between populations. Asterix indicates p-val < 0.05.

Clustering with sNMF shows substantial structure among Madagascar turtle dove populations, with K=2 best explaining the structure in the data (Figure S3). At K=2, individuals from Mauritius and Reunion belong to distinct ancestral populations, while those from Madagascar show equal ancestry from both ancestral populations (Figure 3A). According to this model, individuals 205E and 225E from Mauritius have the highest proportion of mixed ancestry amongst the volcanic island populations. We include K=3 because the PCA indicates three clusters. While individuals from Reunion share the same ancestry, Mauritian individuals exhibit mixed ancestry, consistent with the larger variability observed in the PCA analysis (Figure 3B). Some Mauritian individuals cluster closer to Madagascar than to others from the same population. A small proportion of shared ancestry exists between Reunion and Mauritius. Madagascar individuals remain admixed between two ancestral populations, with one characterizing the current population on Reunion, and the other more frequent in the current population of Mauritius.

**Figure 3.**
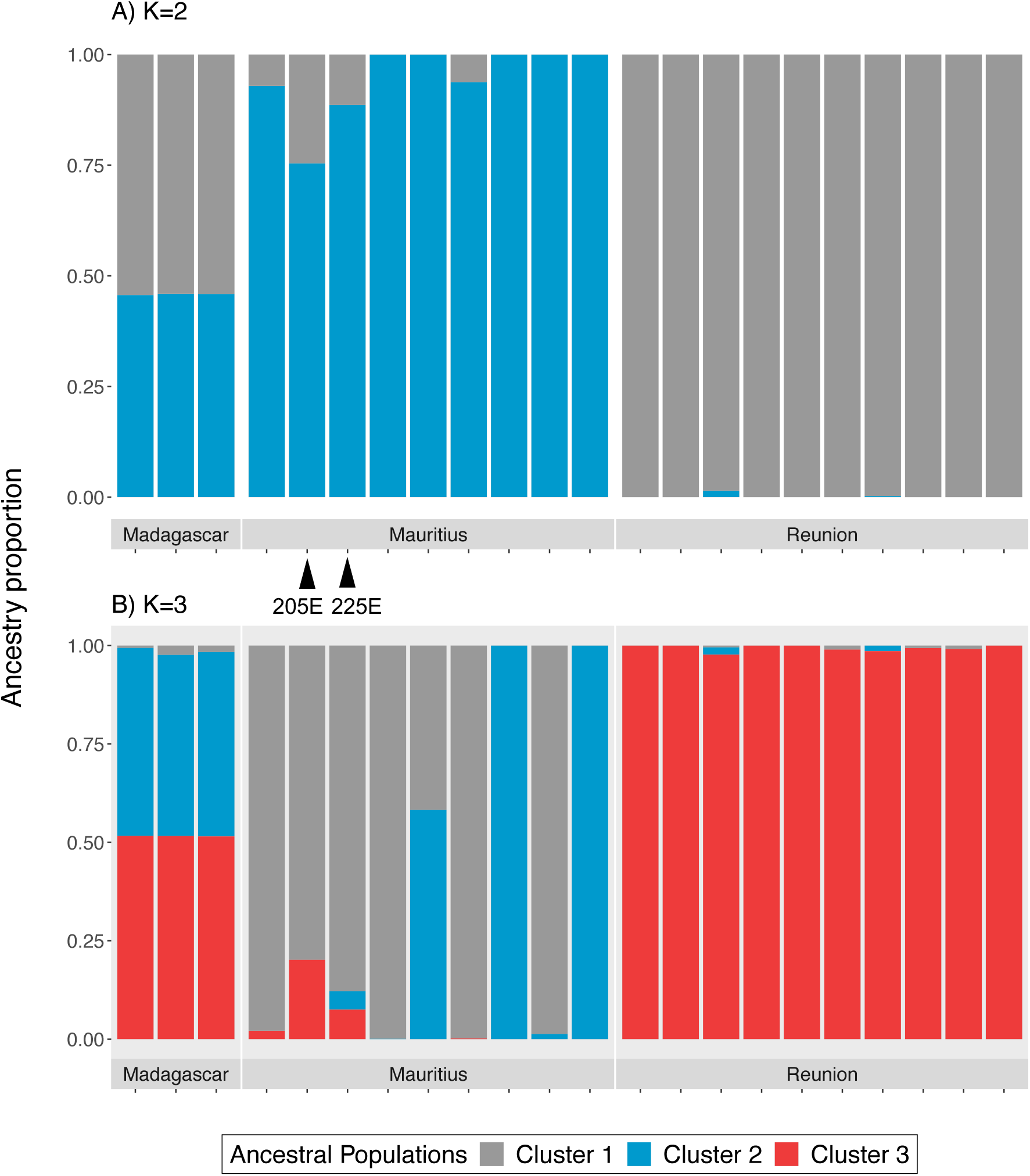
sNMF clustering of *N. picturata* genomes for **(A)** K=2 (corresponding to the lowest cross-entropy) and **(B)** K=3. Labels identify individual samples 205E and 225E.

#### Subfossil samples

PCAs were performed using data from subfossils and modern samples (Figure 4). Sub1 and Sub13, described as *N. cicur or N. picturata* and *N. picturata* respectively (Table S1), cluster with the modern Mauritian lineage. Pairwise IBS distances between these subfossils and modern individuals also indicate high genetic similarity to the Mauritian population (Figures S4 and S5). Sub3 and Sub4, both morphologically identified as either *N. cicur* or *N. picturata* (Table S1), cluster closer to the modern Madagascar lineage. Pairwise IBS distances reveal that Sub3 and Sub4 are genetically distinct from all modern lineages and do not reflect reduced distances to Madagascar individuals (Figures S6 and S7).

**Figure 4.**
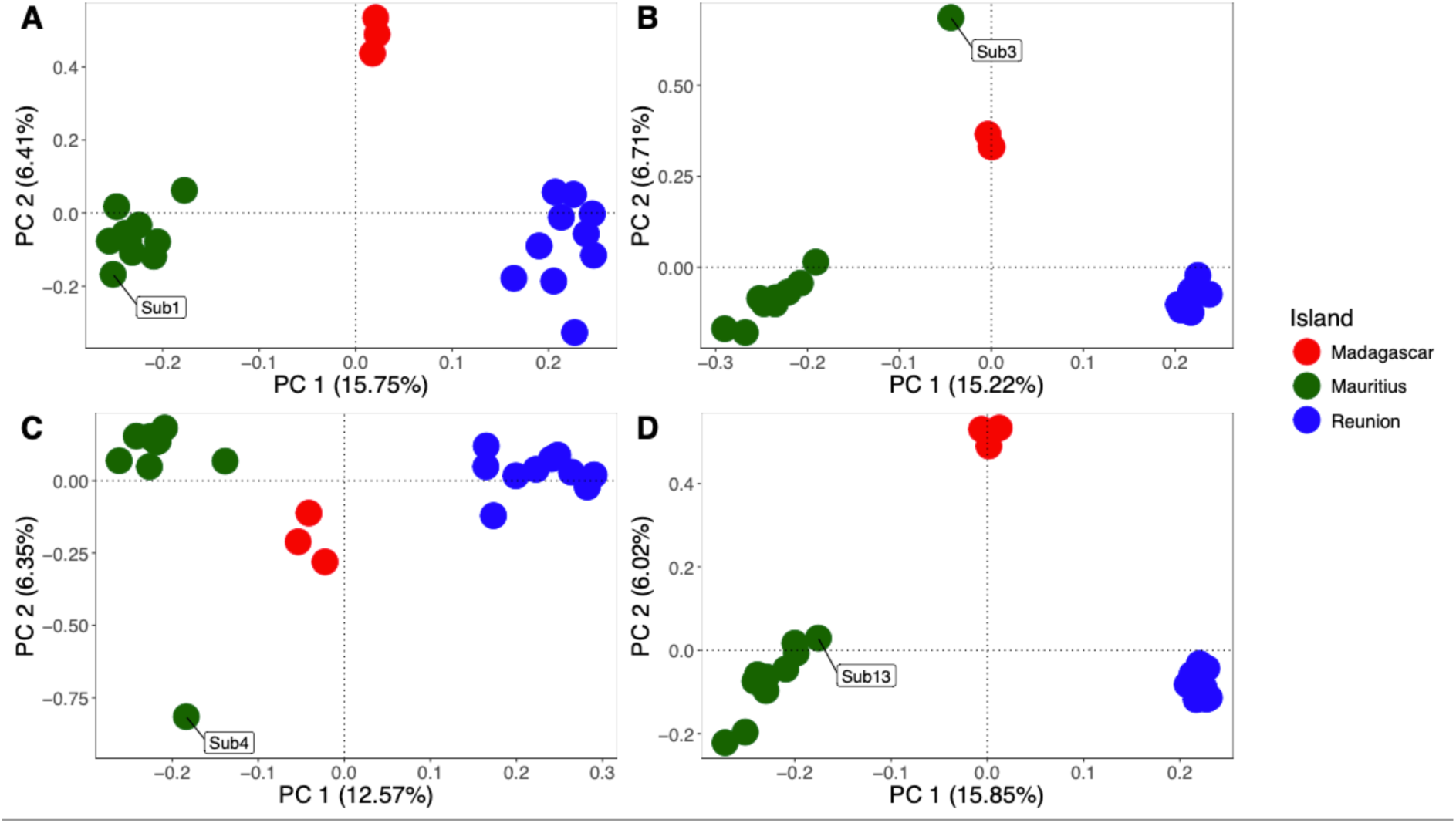
PCA of each subfossil aligned to the modern dataset. Colors represent geographic origin of samples. Subfossils are labelled in plots by their sample name. **(A)** Sub1. **(B)** Sub3. **(C)** Sub4. **(D)** Sub13.

### Genetic diversity and ROH

Genome-wide estimates of 𝜃_π_and 𝜃_w_, indicate that genetic diversity is higher in Madagascar than in Mauritius and Reunion (Table 1), with non-overlapping distributions of scaffold-wide diversity (Figure S8). Volcanic island populations have similar levels of diversity. Positive values of Tajima’s D in volcanic island populations illustrate a modest overrepresentation of high-frequency variants compared to a neutral constant demographic model (Table 1). Conversely, a negative value of Tajima’s D in Madagascar reflects similarly modest overrepresentation of low frequency variants over that expected under neutrality.

**Table 1.** Genetic diversity statistics for *N. picturata* populations.

|  | **Mauritius** | **Reunion** | **Madagascar** |
| --- | --- | --- | --- |
| N | 9 | 10 | 3 |
| $n_{\text{SNP}}$ | 12,112,726 | 12,739,591 | 10,897,013 |
| $\theta\pi$ | $6.0 \times 10^{-3}$ | $6.1 \times 10^{-3}$ | $7.2 \times 10^{-3}$ |
| $\theta W$ | $5.5 \times 10^{-3}$ | $5.6 \times 10^{-3}$ | $7.5 \times 10^{-3}$ |
| Tajima's D | 0.37 | 0.33 | -0.25 |

By calculating the average individual genome-wide H_O_, we find lower genetic variation in the small island populations, with averages of 5.74 and 5.68 heterozygotes per kb (het/kb) for Mauritius and Reunion, respectively (Figure 5). In contrast, the Madagascar population exhibited a higher average of 6.82 het/kb. The variance in average genome-wide H_O_ is greater for the volcanic island populations, especially in Mauritius where it ranges between 5.42 and 5.88 het/kb.

**Figure 5.**
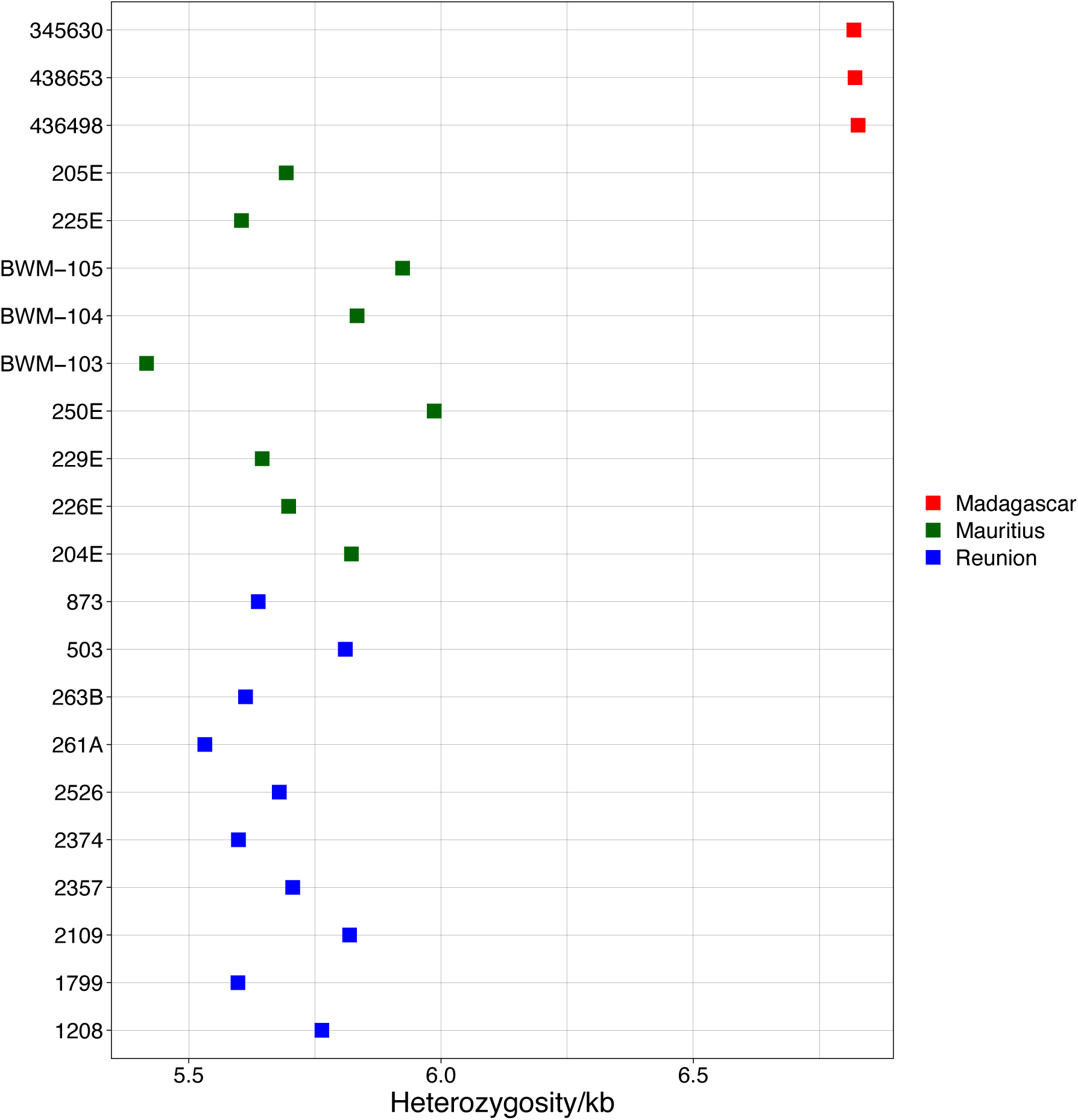
Genome-wide observed heterozygosity by sample. Points are coloured by population.

The lengths of all identified ROH range from 300 kb to 12.5 Mb. Studies consider long ROH to be over 1 Mb (Martin et al., 2023). In the Madagascar population, there are only 5 ROH over 1 Mb, with 4 found in one individual (436498_S44). The longest ROH are in individuals from Mauritius and Reunion (Figure 6). The sum (SROH) and number (NROH) of ROH in each individual are highly correlated (Spearman ρ=0.96, p-val=3.8x10^-6^). Individuals from volcanic islands show greater SROH and NROH than individuals from Madagascar, with the exception of two individuals from Mauritius. Individuals 205E and 225E have ROH comparable to those of Madagascar individuals. Although these individuals have levels of genome-wide H_O_ comparable to the rest of the Mauritian population, examining H_O_ across 300k bp windows of scaffold 1 (the longest scaffold) shows that they have a higher first quartile of heterozygous position per 300k bp windows compared to the other individuals from the volcanic islands (Figure S9).

**Figure 6.**
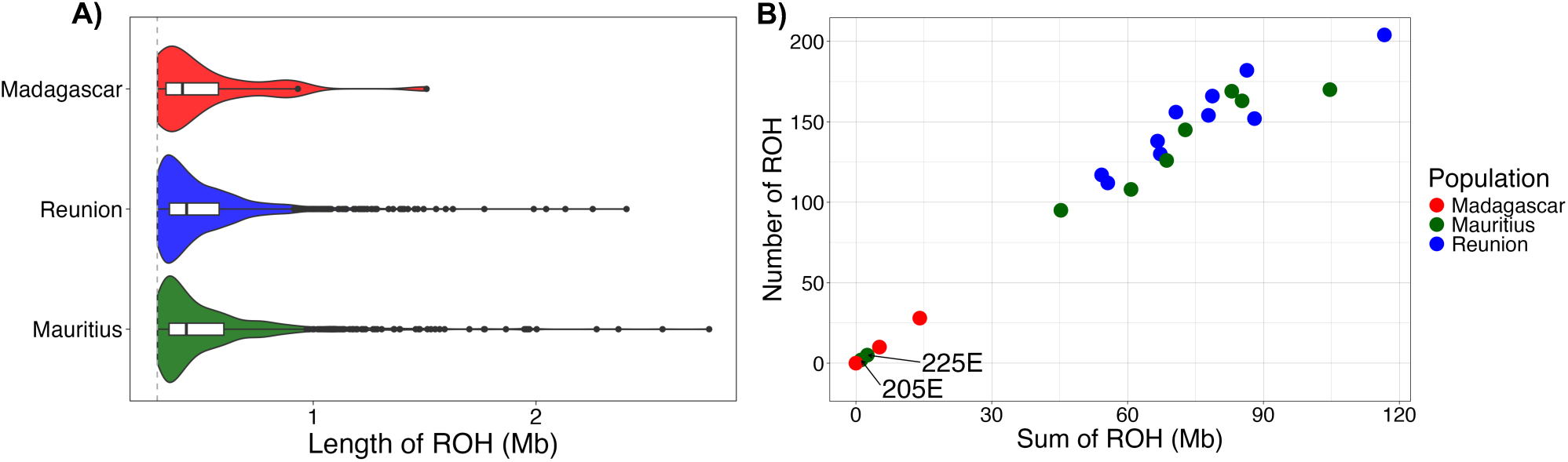
Representation of the length and number of ROH. **(A)** Kernel density plots showing the distribution of ROH length grouped by population. White rectangles represent the interquartile range with the bar as the median. **(B)** Number of ROH (NROH) compared to the sum of the length of ROH (SROH) by individual. Points are coloured by population. Labels identify individual samples 205E and 225E.

### Demographic inferences

PSMC and SMC++ do not yield the same demographic trajectory curves (Figure 7), which we scaled by the mutation rate to provide an approximate time frame. Dynamics of the IICR in the volcanic island populations are similar in their timing and magnitude. PSMC suggests a similar but not overlapping IICR trajectory of individuals from the three islands, while SMC++ clearly suggests a split of Madagascar from Mauritius and Reunion at least 100k YBP. If interpreted in terms of *Ne* variation, inferences of the IICR with SMC++ show drastic declines in Mauritius and Reunion occurring between 1k and 10k YBP. The declines are followed by rapid expansions leading to constant phases. Inferences of the Madagascar population show a more stable dynamic. PSMC inferences, based on single genomes, highlight two individuals from Mauritius (205E and 225E) with differing trends. These two individuals can also be identified through divergent patterns in the sNMF and ROH analyses, but they do not segregate from their own sampling group in intra-population structure analyses (Figure S10).

**Figure 7.**
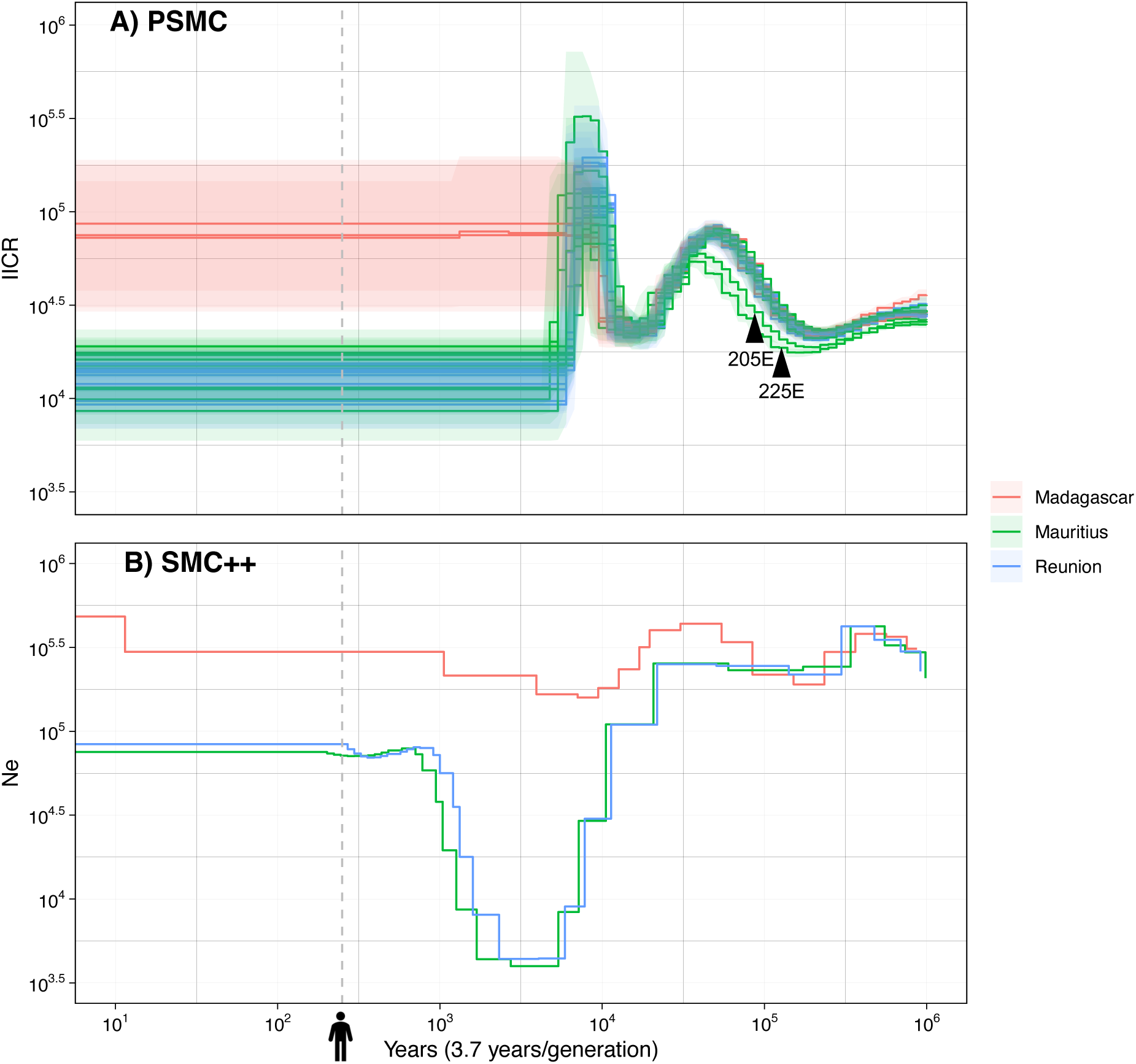
Variation of the IICR/Ne through time in years inferred using **(A)** PSMC. **(B)** SMC++. Dashed grey lines approximately reflect human settlement in the Mascarenes. Colours represent the population of origin. Labels identify individual samples 205E and 225E.

### Demographic modelling

We tested eight demographic scenarios using *fastsimcoal2*. We started by testing tree topology and colonization sequence with MOD1 to MOD6 (Figure S1). MOD2 emerged as the most likely model (Table S3). This model involves two independent colonizations of the Mascarenes from Madagascar in which the sequence is fixed, the first being to Mauritius and the second to Reunion. The estimated colonization times for both islands are similar, at 119,036 and 110,612 YBP, respectively. When comparing MOD2 to MOD6, which also posits two independent colonizations but without a fixed sequence, we find that their likelihood distributions overlap (Figure S11). Given the similarity in colonization times despite the fixed sequence in MOD2 and the negligible difference in likelihoods between MOD2 and MOD6, we retained MOD6 as the most likely tree topology, as it allows for independent estimations of the colonization events. Next, we tested for gene flow by adding asymmetric gene flow to the model in MOD6_MIG. As expected, allowing for gene flow significantly increases the likelihood of the model and permits a lower AIC (Figure S11). Finally, we increase the complexity of the model in MOD6_MIG_TR to account for: i) the variation in coalescence rate inferred for both Mauritius and Reunion using SMC++, which is old enough in terms of generations to exclude a recent time bias (Figure 7); ii) the historical evidence that recent human-mediated translocations of individuals from Madagascar to Mauritius and Reunion were common (Cheke & Hume, 2008; Probst, 2002). MOD6_MIG_TR has the lowest AIC, and a significantly higher likelihood than that of any other model (Table S3).

We ran model MOD6_MIG_TR 200 times and analyzed the resulting parameter distributions. This analysis reveals two distinct scenarios: some runs estimate large current deme sizes for Mauritius and Reunion (N_MAU_ and N_REU_) with low translocation rates from Madagascar (TR_MAD→MAU_ and TR_MAD→REU_), while others suggest smaller deme sizes with higher translocation rates (Figure S12 and S13). Thus, while some runs indicate large current population sizes, others suggest the occurrence of a significant and recent bottleneck. Runs estimating larger deme sizes and lower translocation rates tend to have higher likelihoods, a finding that diverges from historical records indicating frequent translocations from Madagascar to Mauritius and Reunion, as well as recent population declines (Cheke & Hume, 2008; Probst, 2002). Instead of focusing on the run with the highest likelihood, we present two runs that exemplify the contrasting relationships between current deme size and translocation rates, that we name H_N_L_T_ and L_N_H_T_ (Figure 8).

**Figure 8.**
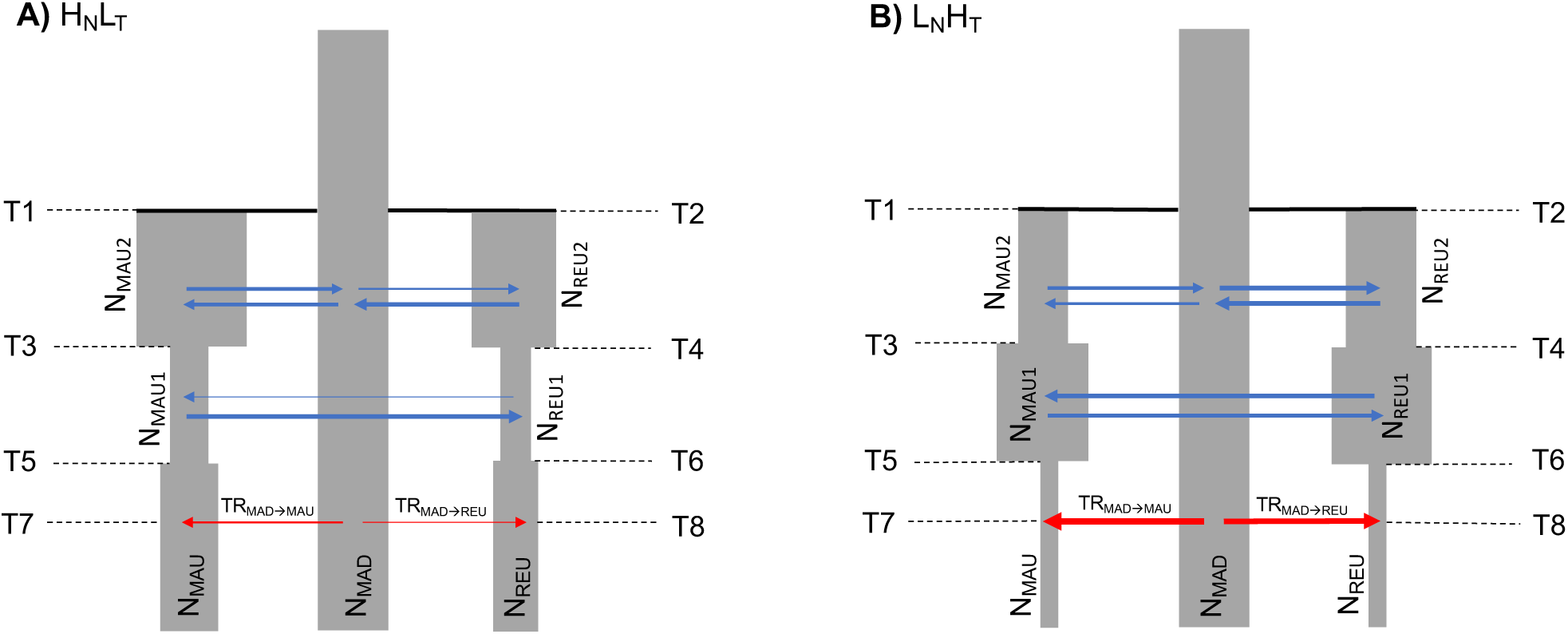
Representation of the most likely demographic scenario (MOD6_MIG_TR) scaled to parameter estimates of **(A)** H_N_L_T_. **(B)** L_N_H_T_.

In H_N_L_T_, N_MAU_ and N_REU_ are estimated at 102,403 and 43,602, respectively, whereas in L_N_H_T_, these estimates decrease to 540 and 629 (Table 2). The translocation rates from Madagascar to Mauritius and Reunion (TR_MAD→MAU_ and TR_MAD→REU_) are estimated at 0.02 and 0.004 in H_N_L_T_ compared to 0.19 and 0.12 in L_N_H_T_. The patterns of *Ne* variation from colonization to the present differ notably between H_N_L_T_ and L_N_H_T_. In H_N_L_T_, *Ne* decreases after colonization at times T3 and T4 and increases in the more recent past. Conversely, in L_N_H_T_, *Ne* increases at T3 and T4 but subsequently dramatically declines in recent times. These recent changes in *Ne* occur at time points T5 and T6, estimated in H_N_L_T_ at 207 and 366 YBP and in L_N_H_T_ at 307 and 370 YBP for Mauritius and Reunion respectively, periods that align with human settlement in the Mascarenes.

**Table 2.** ML values of demographic parameters estimated under H_N_L_T_ and L_N_H_T_ of MOD6_MIG_TR and their 5% and 95% confidence intervals computed after parametric bootstrap. Times are expressed in years (assuming a generation time of 3.7 years). Migration rates are indicated in a forward direction.

| Parameter | $H_N L_T$ ML point estimates | $L_N H_T$ ML point estimates |
| --- | --- | --- |
| $N_{MAU}$ | 102,403 (11,229-338,705) | 540 (530-1,717) |
| $N_{REU}$ | 43,602 (763-402,348) | 629 (410-1,740) |
| $N_{MAD}$ | 479,042 (402,348-464,609) | 512,869 (387,110-761,306) |
| $N_{MAU1}$ | 43,227 (47,867-143,853) | 532,314 (137,427-764,464) |
| $N_{REU1}$ | 26,150 (13,443-402,894) | 617,752 (166,258-767,980) |
| $N_{MAU2}$ | 1,394,066 (450,975-976,293) | 53,309 (38,823-472,393) |
| $N_{REU2}$ | 608,445 (263,290-937,942) | 178,437 (51,034-457,010) |
| T1 | 659,514 (687,175-1,246,666) | 1,261,478 (947,048-2,797,144) |
| T2 | 698,623 (570,447-1,437,080) | 1,449,494 (939,914-2,750,711) |
| T3 | 34,787 (30,843-99,396) | 836,167 (38,427-1,325,836) |
| T4 | 22,718 (9,705-534,856) | 736,996 (37,933-1,386,905) |
| T5 | 207 (114-359) | 307 (243-705) |
| T6 | 366 (122-650) | 370 (247-715) |
| T7 | 170 (77-322) | 270 (206-668) |
| T8 | 296 (85-613) | 333 (210-678) |
| $m_{REU \rightarrow MAU}$ | $1.88 \times 10^{-6}$ ($2.97 \times 10^{-7}$-$5.12 \times 10^{-6}$) | $7.42 \times 10^{-5}$ ($8.04 \times 10^{-6}$-$7.94 \times 10^{-5}$) |
| $m_{MAD \rightarrow MAU}$ | $1.33 \times 10^{-5}$ ($2.13 \times 10^{-6}$-$1.89 \times 10^{-5}$) | $1.56 \times 10^{-5}$ ($3.19 \times 10^{-6}$-$5.65 \times 10^{-5}$) |
| $m_{MAU \rightarrow REU}$ | $4.17 \times 10^{-5}$ ($2.57 \times 10^{-5}$-$5.64 \times 10^{-5}$) | $6.67 \times 10^{-5}$ ($3.92 \times 10^{-6}$-$8.77 \times 10^{-5}$) |
| $m_{MAD \rightarrow REU}$ | $4.70 \times 10^{-6}$ ($6.28 \times 10^{-7}$-$2.94 \times 10^{-5}$) | $7.79 \times 10^{-5}$ ($9.59 \times 10^{-7}$-$5.79 \times 10^{-5}$) |
| $m_{MAU \rightarrow MAD}$ | $1.44 \times 10^{-5}$ ($1.18 \times 10^{-6}$-$1.16 \times 10^{-5}$) | $1.96 \times 10^{-5}$ ($2.65 \times 10^{-6}$-$7.32 \times 10^{-5}$) |
| $m_{REU \rightarrow MAD}$ | $3.03 \times 10^{-5}$ ($2.15 \times 10^{-5}$-$5.19 \times 10^{-5}$) | $8.28 \times 10^{-5}$ ($1.42 \times 10^{-6}$-$7.44 \times 10^{-5}$) |
| $TR_{MAD \rightarrow MAU}$ | 0.02 (0.001-0.02) | 0.19 (0.02-0.33) |
| $TR_{MAD \rightarrow REU}$ | 0.004 (0.001-0.1) | 0.12 (0.02-0.55) |

Some parameter estimations remain similar across runs of MOD6_MIG_TR. Despite large confidence intervals, the model consistently supports the two independent colonization events both occurring long before human arrival in the Mascarenes. In both runs, Mauritius is estimated to have been colonized after Reunion, with T1 estimated at 659,514 YBP (687,175-1,246,666 YBP) and 1,261,478 YBP (947,048-2,797,144 YBP) in H_N_L_T_ and L_N_H_T_, respectively (Table 2). The colonization of Reunion is estimated at 698,623 YBP (570,447-1,437,080 YBP) in H_N_L_T_ and 1,449,494 YBP (939,914-2,750,711 YBP) in L_N_H_T_. Both runs also indicate gene flow with the migration rates ranging from 1.88x10^-6^ (m_REU→MAU_) to 4.17x10^-5^ (m_MAU→REU_) in H_N_L_T_ and from 1.56x10^-5^ (m_MAD→MAU_) to 8.28x10^-5^ (m_REU→MAD_) in L_N_H_T_. In both runs, the estimates of translocation times correspond to a period of human establishment in the Mascarenes. The translocation from Madagascar to Mauritius (T7) is estimated to have occurred 170 YBP in H_N_L_T_ and 270 YBP ago in L_N_H_T_. Similarly, translocation from Madagascar to Reunion (T8) is estimated at 296 YBP in H_N_L_T_ and 333 YBP in L_N_H_T_ (Table 2).

### Goodness-of-fit of the most likely model

#### Simulated structure and diversity

To determine the most plausible of the two MOD6_MIG_TR scenarios, we examined the goodness-of-fit of H_N_L_T_ and L_N_H_T_ by simulating a 10,934,710 bp scaffold (the length of scaffold 1) using the ML parameter estimates obtained from the 100 parametric bootstraps of each run. Both runs accurately reconstruct the observed population structure. PCA from simulations of H_N_L_T_ and L_N_H_T_ consistently reveal three population clusters (Figure S14). Concordant with the observed pairwise Fst values (Figure 2), the highest estimated divergence is between Mauritius and Reunion (0.15 in both runs; Figure S15). sNMF clusters are almost identical between H_N_L_T_ and L_N_H_T_ (Figure S16). As in the observed data, the populations do not cluster in separate groups when K=3. Both runs successfully replicate the observed genetic diversity (Figure S17). Madagascar consistently shows higher genetic diversity than the volcanic islands. Minor differences between the observed and simulated genomic average of summary statistics, particularly in Tajima’s D, can be attributed to the inherent variation observed both between and within scaffolds (Figure S8; and specifically for scaffold 1, Figure S18) and most likely reflecting genomic complexity (which can be hardly reproduced with simulations).

#### IICR and LD statistics

We further simulated the IICR from parametric bootstraps of H_N_L_T_ and L_N_H_T_ (Figure 9). The Madagascar IICR remains relatively stable and elevated across runs. In contrast, IICR trajectories for Mauritius and Reunion show significant differences between the two runs. In H_N_L_T_, the IICR suggests stable population sizes throughout the islands’ histories, with the exception of a single ancestral shift around 10k YBP. In L_N_H_T_, the IICR profiles for Mauritius and Reunion reveal pronounced bottlenecks approximately 500 YBP, leading to reduced current population sizes. The IICR from L_N_H_T_ show larger variance in recent times in Mauritius and Reunion compared to H_N_L_T_, consistent with the variance estimated in the bottleneck intensity (Figure S19). When we re-analyzed simulated data from H_N_L_T_ and L_N_H_T_ using PSMC and SMC++ we obtained similar trajectories, but the recent bottleneck in L_N_H_T_ was not retrieved by any of the two algorithms (Figure S20).

**Figure 9.**
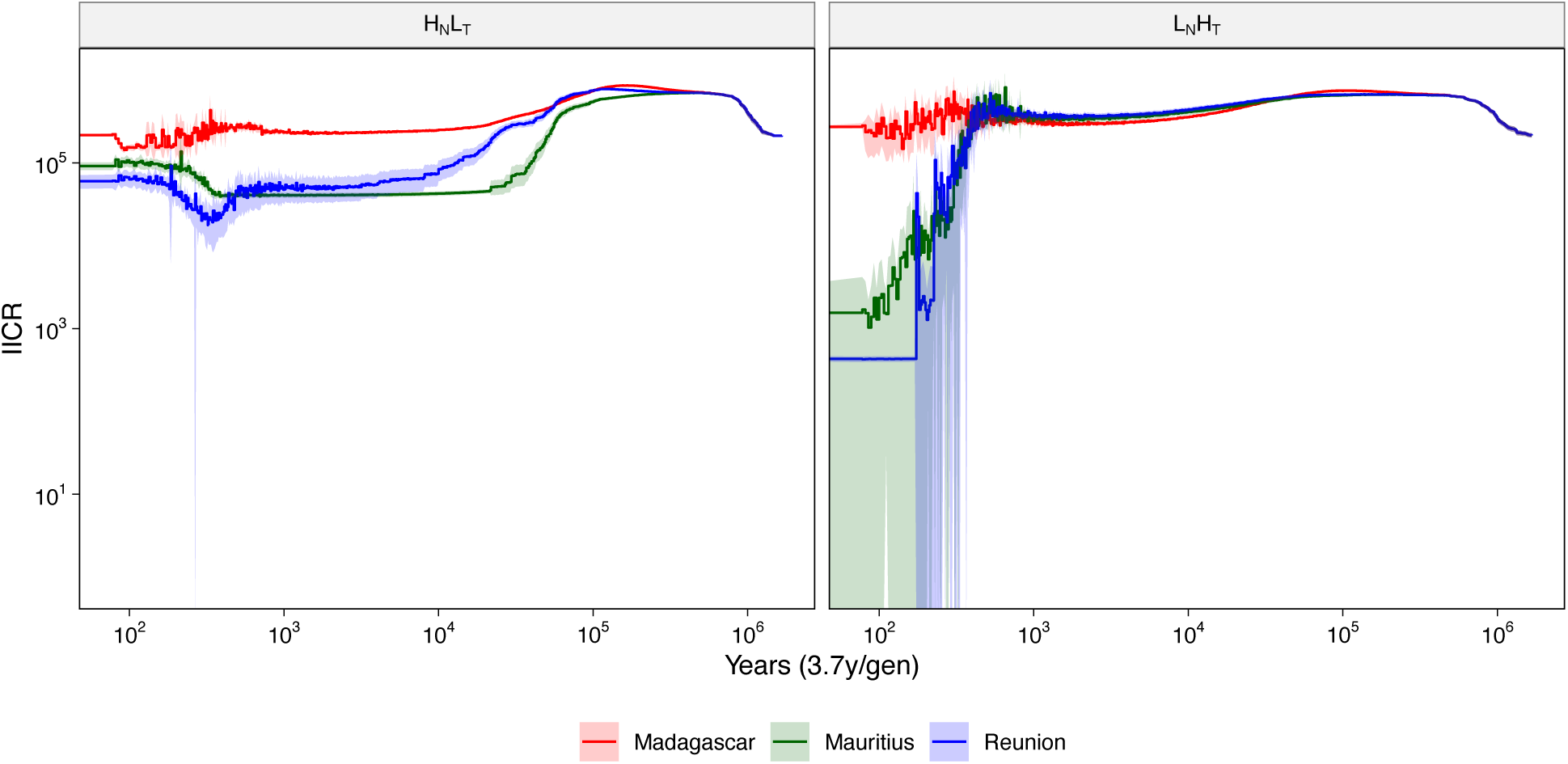
Mean simulated IICR (IICRsim) and its 95% CI over time computed from parametric bootstraps of: **(A)** H_N_L_T_. **(B)** L_N_H_T_.

To explore additional genomic information that was not part of the inferential process leading to the selection of the most likely scenario (MOD6_MIG_TR), we simulated the theoretical expectations of LD decay using parameters from parametric bootstraps of H_N_L_T_ and L_N_H_T_. As expected, both theoretical and observed LD decrease with increasing recombination distances, but the decay is different between the two runs (Figure 10). Simulations from L_N_H_T_ predict higher LD levels at higher recombination distances compared to H_N_L_T_. At these higher recombination distances, the confidence intervals around the observed LD, as measured by 𝔼[D_z_], partially overlap with the theoretical LD from L_N_H_T_, but not from H_N_L_T_. Generally speaking, the decay observed in the real data is lower decays simulated from the two runs, but closer to L_N_H_T_.

**Figure 10.**
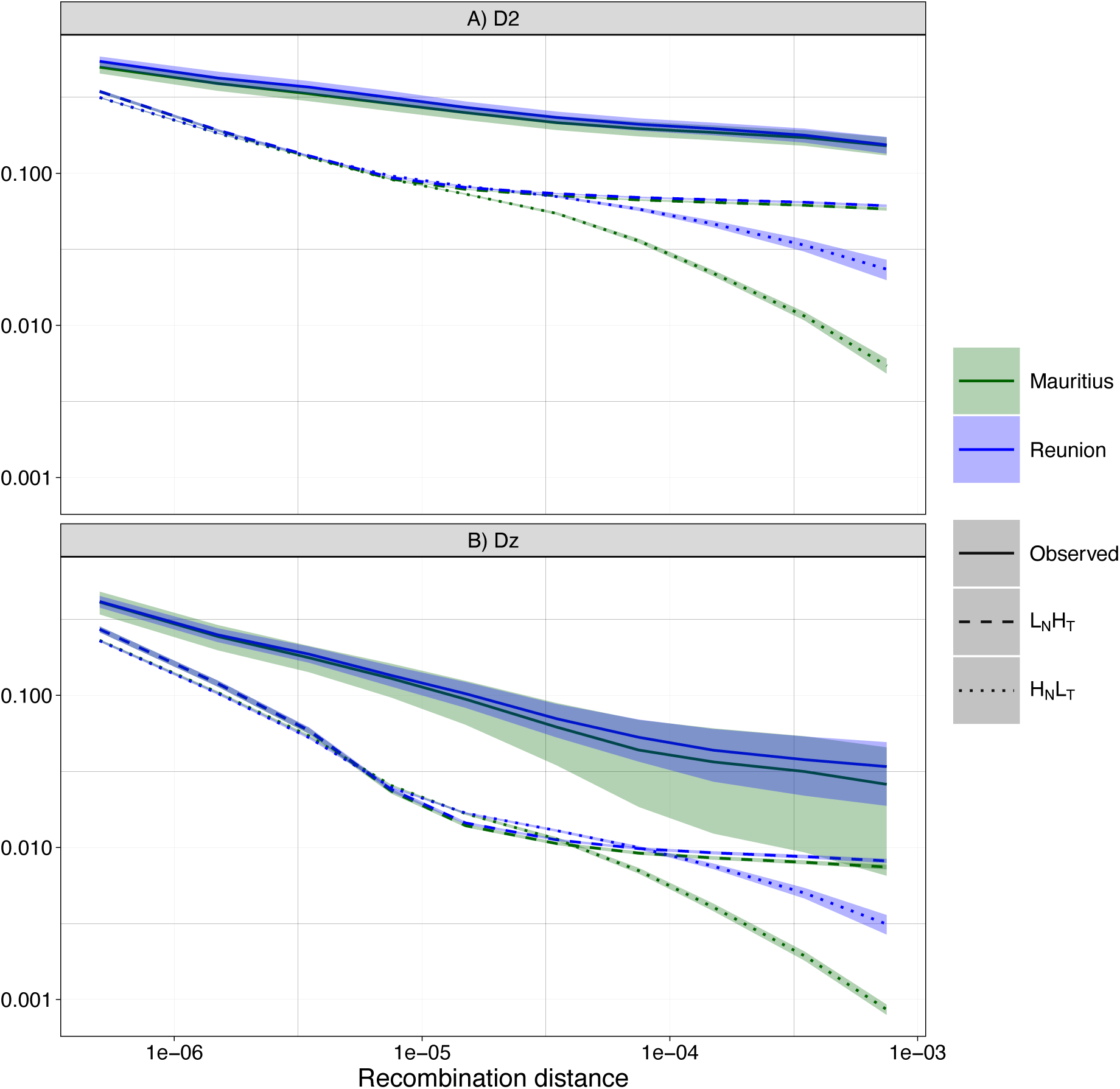
Theoretical expectations of LD decay by recombination distance in comparison to observed LD decay computed with *moments.LD*. Theoretical expectations are computed from parametric bootstraps of H_N_L_T_ and L_N_H_T_. LD is estimated by: **(A)** 𝔼[D^2^]. **(B)** 𝔼[D_z_].

## Discussion

The spread of non-native species is of global conservation concern (Simberloff et al., 2013). Designing effective conservation strategies to manage non-native species is often hindered by a lack of clarity on species’ status as native or non-native. Our study takes advantage of the known recent timing of first human arrival in the Mascarenes 400 years ago (A. Cheke & Hume, 2008), along with sequencing of modern and ancient DNA, to examine the status of an avian species of concern for conservation programs in the Mascarenes. We used the dataset to analyze genetic structure and diversity, estimate ROH, and infer demographic history, presenting a likely scenario of colonization of the Mascarenes by the Madagascar turtle dove. We further use a comprehensive population genetic approach to resolve ambiguities in demographic modelling. Our key finding, with important conservation implications, is that *N. picturata* is native to both Mauritius and Reunion. Two independent lines of evidence demonstrate that *N. picturata* was established in the Mascarenes long before human arrival, indicating that its colonization was natural rather than human-mediated.

First, two of the *Nesoenas* subfossils from Mauritius cluster with the modern *N. picturata* population of which one (Sub1) is from the Mare aux Songes horizon radiocarbon dated at 4200 YBP (Hume et al. 2015), demonstrating that the extant Mauritius population is at least 4200 years old, predating human arrival. We also identified subfossils showing intermediate divergence between the current populations of Mauritius and Madagascar in PCA. Genetic distance calculations reveal that these subfossils are genetically distinct from all modern samples. The divergence of certain subfossils from modern populations could represent an extinct, divergent lineage, such as *N. cicur*, or it may result from insufficient confidence in genotypes due to low genomic coverage, or age-related biases. As expected, obtaining genome-wide data from only 30-50 mg of subfossil bone originating from low preservation quality tropical environments (swamp, cave and boulder scree) was challenging. The biggest challenge encountered was that despite optimization of the temperature used for hybridization, we obtained low capture specificity and therefore low overlap between samples. This appears to be a hard constraint of the ancient DNA itself. Nonetheless, the capture was successful in enhancing endogenous DNA yield by up to five-fold, and obtaining up to 680,000 variants between modern and ancient samples. As such, our study shows that ancient DNA can confirm identity of populations of non-model colonizing species on timescales of 100-4000 years, spanning and exceeding those in which human interventions have taken place, and yet in which demographic inferences are generally challenging.

Second, despite the challenges of demographic inferences from modern DNA on timescales of up to 4000 years, demographic inference proves paramount in resolving the timing of the colonizations giving rise to the living populations of Mauritius and Reunion on longer timescales. Multiple runs of the most likely coalescent-based model consistently estimate colonization times for Mauritius and Reunion that predate human arrival, even when accounting for confidence intervals (Table 2). In H_N_L_T_, colonization events are estimated to have occurred hundreds of thousands of YBP, while L_N_H_T_ estimates approach one million YBP. Additionally, the inferred coalescent rate trajectories (particularly from SMC++), indicate that the volcanic island populations diverged from Madagascar long before first human arrival in the Mascarenes (Figure 7). The inferred IICR depends on the underlying demographic history and can reflect population divergence (Lesturgie et al., 2022).

Our results also demonstrate the nature of the colonization events, indicating that Mauritius and Reunion were independently colonized from Madagascar. The PCA and Fst values reveal greater genetic divergence between the two volcanic islands (Fst = 0.15), separated by 164 km, than between either island and Madagascar, a distance of ≥ 665 km (Thébaud et al., 2009; Figure 2). The lack of correlation between genetic and geographic distances contrasts with certain other studies of birds colonizing multiple islands (e.g. Spurgin et al., 2014). Furthermore, differentiation even between distant islands is often weak in birds (e.g. Fst of 0.04 in Berthelot’s pipit between islands separated by 470 km; Martin et al., 2021). The high differentiation we uncover among volcanic island populations and intermediate position of the Madagascar population in structure analyses support independent colonization events from Madagascar. It is unlikely that an admixed Madagascar population biased this inference, as a model in which the Madagascar population resulted from admixture between Mauritius and Reunion populations gains less support than other models. In contrast, models of independent colonization show significantly better AIC scores.

In testing scenarios of colonization history for *N. picturata*, this study highlights the necessity of using an integrated approach to validate the biological relevance of demographic models. Although the coalescent framework facilitates the comparison of complex scenarios, it does not guarantee goodness of fit, making posterior validation essential. We identified a likely model by maximizing the resemblance between observed and predicted SFS. Interestingly, this model yields two distinct sets of parameter estimates for current deme sizes and translocation rates on the two volcanic islands. H_N_L_T_ suggests large current deme sizes with negligible translocation rates from Madagascar. Conversely, L_N_H_T_ indicates significant bottlenecks, with strengths of approximately 950x linked to human settlement in the Mascarenes and associated with high translocation rates (Table 2). Despite a higher likelihood, H_N_L_T_ is not consistent with historical reports concerning introductions and population declines. To identify the most biologically relevant scenario, we assessed goodness-of-fit using additional summary statistics, such as IICR and LD metrics. Simulations from the ML parameter values of both runs H_N_L_T_ and L_N_H_T_ successfully replicate the observed genetic structure and diversity in Mauritius and Reunion (Figures S11-S14). In contrast, simulations of the IICR over time reveal distinct demographic trajectories, with L_N_H_T_ showing recent changes consistent with bottlenecks. Inferring the IICR from observed data may not reveal such changes as sequential Markovian approaches can fail to detect recent variation in coalescence rate, especially when genetic diversity is large (Palsbøll et al., 2013). Indeed, PSMC and SMC++ analyses of data simulated under L_N_H_T_ do not capture the recent bottleneck (Figure S20). We also examined patterns of LD decay, which has potential to resolve more recent events since recombination occurs more frequently than mutations (Boitard et al., 2016). By analyzing theoretical and observed LD decay patterns, we found that these patterns differentiate H_N_L_T_ from L_N_H_T_, and that L_N_H_T_ better predicts the observed LD decay at higher recombination distances. Overall, while neither run captures all aspects of the observed data, L_N_H_T_ better aligns with the observed LD pattern. Moreover, SFS-based statistics (Tajima’s D and genetic diversity) and the IICR do not effectively differentiate between runs as SFS statistics behave similarly and IICR inferences lack sensitivity to recent variations.

Our study also illustrates why relying solely on genetic diversity metrics for conservation can be misleading. Indeed, the genetic diversity observed in Mauritius and Reunion is exceptional for colonizing populations (Wittwer et al., 2023). This diversity is reflected in the genome-wide H_O_, that ranges from 5 to 7 het/kb (Figure 5), which is higher than many other species that have undergone recent population contraction (e.g. 0.5-2 het/kb for fin whales; Nigenda-Morales et al., 2023). Although such high levels of genetic diversity might indicate large and stable populations, our integrated approach demonstrates that, in this case, the observed diversity is more likely shaped by the translocation events identified in L_N_H_T_. Studies of recently bottlenecked populations have shown that the immigration of a few individuals can significantly increase genetic diversity (Åkesson et al., 2016; Consuegra et al., 2005; Mills & Allendorf, 1996). Further evidence of these translocation events is provided by our recombination-based demographic analyses (PSMC) and ROH estimations, which reveal that two individuals from Mauritius (205E and 225E) exhibit genetic patterns distinct from others in their population (Figure 6 and 7) that cannot be attributed to low sequence depths (Table S1). These individuals have lower SROH and NROH (close to those from Madagascar; Figure 6), consistent with them being the descendants of admixture between lineages, which breaks up ROH (Ceballos et al., 2018; Foote et al., 2021; Pemberton et al., 2012). Clustering analysis with K=2 (which has the lowest cross entropy) shows that the anomalous individuals have the highest mixed ancestry proportion in the Mauritian population, suggesting that they descend from recent admixture between the Madagascar and Mauritius lineages. More generally, these results reflect the significant impact that demographic processes, such as translocations, can have on shaping genetic diversity.

In conclusion, our findings support revising the status of *N. picturata* as native to both Mauritius and Reunion. Understanding the role of past human intervention in the distribution of a species is an important problem in conservation at a global scale (Gallardo et al., 2015). Until the last decade, a major challenge in determing the role of human intervention was the difficulty of resolving recent demographic history with genetic data. Methods employing LD to infer recent demographic history (Fournier et al., 2023; Ragsdale & Gravel, 2019; Santiago et al., 2020) present significant prospects, despite being relatively new. Our study illustrates how data from subfossils can be used to complement and corroborate demographic inferences from whole genome modeling on recent time scales. We find that the two independent lines of inference concur, strengthening evidence for pre-human colonization. Our study also highlights the necessity of rigorous *a posteriori* validation using summary statistics independent of the inferential process to minimize the acceptance of biologically irrelevant models. Going forward, conservation genetic studies would benefit from integrating multiple summary statistics into a unified analysis.

## Acknowledgments

We are grateful to the SEOR for providing samples from Reunion and for their involvement in this project. We thank the Mauritius National Parks and Conservation Service and the Mauritian Wildlife Foundation for their valuable assistance and for facilitating our fieldwork. We also thank Céline Bon and Angelika Borowiecka for their advice relating to bioinformatic pipelines. We are appreciative of the Genotoul bioinformatics platform (Bioinfo Genotoul; http://bioinfo.genotoul.fr/) and the MNHN PCIA cluster for providing computing resources. This research was supported by the Agence Nationale de la Recherche ANR-20-E02-0009 (Suscept-Ext).

High-throughput PacBio sequencing was performed by the ICGex NGS platform of the Institut Curie supported by the grants ANR-10-EQPX-03 (Equipex) and ANR-10-INBS-09-08 (France Génomique Consortium) from the Agence Nationale de la Recherche (“Investissements d’Avenir” program), by the ITMO-Cancer Aviesan (Plan Cancer III) and by the SiRIC-Curie program (SiRIC Grant INCa-DGOS-465 and INCa-DGOS-Inserm_12554). PacBio data management, quality control and primary analysis were performed by the Bioinformatics platform of the Institut Curie.

## Data Accessibility and Benefit-Sharing Section

### Data Accessibility Statement

This Whole Genome Shotgun project has been deposited at DDBJ/ENA/GenBank under the accession JBJUWA000000000. Raw sequence reads will be deposited in NCBI SRA undern the accession numbers XXXX-XXXX.

### Benefit-Sharing Statement

Benefits Generated: Benefits from this research arise from the sharing of our data on public databases as described above. A research collaboration was developed with conservation organisations from the countries from which we obtained genetic samples and our research adresses some of their priority concerns regarding the conversation management of the Madagascar turtle dove.

## References

1. Åkesson, M., Liberg, O., Sand, H., Wabakken, P., Bensch, S., & Flagstad, Ø. (2016). Genetic rescue in a severely inbred wolf population. Molecular Ecology, 25(19), 4745–4756. 10.1111/mec.13797

2. Austin, J. J., Arnold, E. N., & Jones, C. G. (2004). Reconstructing an island radiation using ancient and recent DNA: The extinct and living day geckos (Phelsuma) of the Mascarene islands. Molecular Phylogenetics and Evolution, 31(1), 109–122. 10.1016/j.ympev.2003.07.011

3. Barré, N. (1983). Distribution et abondance des oiseaux terrestres de l’île de la Réunion (Océan Indien). Revue d’Écologie (La Terre et La Vie), 37(1), 37–85. 10.3406/revec.1983.4762

4. Beaumont, M. A. (2010). Approximate Bayesian Computation in Evolution and Ecology. Annual Review of Ecology, Evolution, and Systematics, 41(1), 379–406. 10.1146/annurev-ecolsys-102209-144621

5. Beerli, P. (2006). Comparison of Bayesian and maximum-likelihood inference of population genetic parameters. Bioinformatics, 22(3), 341–345. 10.1093/bioinformatics/bti803

6. Beichman, A. C., Phung, T. N., & Lohmueller, K. E. (2017). Comparison of Single Genome and Allele Frequency Data Reveals Discordant Demographic Histories. 7, 360–3620.

7. Bertelsmeier, C., Bonnamour, A., Garnas, J. R., Liu, T., Perreault, R., & Ollier, S. (2025). Temporal dynamics and global flows of insect invasions in an era of globalization. Nature Reviews Biodiversity. 10.1038/s44358-025-00016-1

8. Boitard, S., Rodríguez, W., Jay, F., Mona, S., & Austerlitz, F. (2016). Inferring Population Size History from Large Samples of Genome-Wide Molecular Data—An Approximate Bayesian Computation Approach. PLOS Genetics, 12(3), e1005877. 10.1371/journal.pgen.1005877

9. Bolger, A. M., Lohse, M., & Usadel, B. (2014). Trimmomatic: A flexible trimmer for Illumina sequence data. Bioinformatics, 30(15), 2114–2120. 10.1093/bioinformatics/btu170

10. Buckley, Y. M., & Catford, J. (2016). Does the biogeographic origin of species matter? Ecological effects of native and non-native species and the use of origin to guide management. Journal of Ecology, 104(1), 4–17. 10.1111/1365-2745.12501

11. Ceballos, F. C., Joshi, P. K., Clark, D. W., Ramsay, M., & Wilson, J. F. (2018). Runs of homozygosity: Windows into population history and trait architecture. Nature Reviews Genetics, 19(4), 220–234. 10.1038/nrg.2017.109

12. Cheke, A. (1987). An ecological history of the Mascarene Islands, with particular reference to extinctions and introductions of land vertebrates. In Studies of Mascarene Island Birds (pp. 5–89). Cambridge University Press.

13. Cheke, A. (2005). Naming segregates from the Columba–Streptopelia pigeons following DNA studies on phylogeny. Bulletin of the British Ornithologists’ Club, 125(4), 293–295.

14. Cheke, A. (2013). Extinct birds of the Mascarenes and Seychelles—A review of the causes of extinction in the light of an important new publication on extinct birds. 21, 4–19.

15. Cheke, A., & Hume, J. (2008). Lost Land of the Dodo. A&C Black Publisher.

16. Consuegra, S., Verspoor, E., Knox, D., & García De Leániz, C. (2005). Asymmetric gene flow and the evolutionary maintenance of genetic diversity in small, peripheral Atlantic salmon populations. Conservation Genetics, 6(5), 823–842. 10.1007/s10592-005-9042-4

17. Danecek, P., Bonfield, J. K., Liddle, J., Marshall, J., Ohan, V., Pollard, M. O., Whitwham, A., Keane, T., McCarthy, S. A., Davies, R. M., & Li, H. (2021). Twelve years of SAMtools and BCFtools. GigaScience, 10(2), giab008. 10.1093/gigascience/giab008

18. DePristo, M. A., Banks, E., Poplin, R., Garimella, K. V., Maguire, J. R., Hartl, C., Philippakis, A. A., del Angel, G., Rivas, M. A., Hanna, M., McKenna, A., Fennell, T. J., Kernytsky, A. M., Sivachenko, A. Y., Cibulskis, K., Gabriel, S. B., Altshuler, D., & Daly, M. J. (2011). A framework for variation discovery and genotyping using next-generation DNA sequencing data. Nature Genetics, 43(5), 491–498. 10.1038/ng.806

19. Estoup, A., Beaumont, M., Sennedot, F., Moritz, C., & Cornuet, J.-M. (2004). Genetic analysis of complex demographic scenarios: Spatially expanding populations of the cane toad, Bufo marinus. Evolution, 58(9), 2021–2036. 10.1111/j.0014-3820.2004.tb00487.x

20. Estoup, A., & Guillemaud, T. (2010). Reconstructing routes of invasion using genetic data: Why, how and so what?: RECONSTRUCTING ROUTES OF INVASION. Molecular Ecology, 19(19), 4113–4130. 10.1111/j.1365-294X.2010.04773.x

21. Excoffier, L., Dupanloup, I., Huerta-Sánchez, E., Sousa, V. C., & Foll, M. (2013). Robust Demographic Inference from Genomic and SNP Data. PLoS Genetics, 9(10), e1003905. 10.1371/journal.pgen.1003905

22. Fages, A., Hanghøj, K., Khan, N., Gaunitz, C., Seguin-Orlando, A., Leonardi, M., McCrory Constantz, C., Gamba, C., Al-Rasheid, K. A. S., Albizuri, S., Alfarhan, A. H., Allentoft, M., Alquraishi, S., Anthony, D., Baimukhanov, N., Barrett, J. H., Bayarsaikhan, J., Benecke, N., Bernáldez-Sánchez, E., … Orlando, L. (2019). Tracking Five Millennia of Horse Management with Extensive Ancient Genome Time Series. Cell, 177(6), 1419–1435.e31. 10.1016/j.cell.2019.03.049.

23. Foote, A. D., Hooper, R., Alexander, A., Baird, R. W., Baker, C. S., Ballance, L., Barlow, J., Brownlow, A., Collins, T., Constantine, R., Dalla Rosa, L., Davison, N. J., Durban, J. W., Esteban, R., Excoffier, L., Martin, S. L. F., Forney, K. A., Gerrodette, T., Gilbert, M. T. P., … Morin, P. A. (2021). Runs of homozygosity in killer whale genomes provide a global record of demographic histories. Molecular Ecology, 30(23), 6162–6177. 10.1111/mec.16137

24. Fournier, R., Tsangalidou, Z., Reich, D., & Palamara, P. F. (2023). Haplotype-based inference of recent effective population size in modern and ancient DNA samples. Nature Communications, 14(1), 7945. 10.1038/s41467-023-43522-6

25. Freed, L. A., & Cann, R. L. (2009). Negative Effects of an Introduced Bird Species on Growth and Survival in a Native Bird Community. Current Biology, 19(20), 1736–1740. 10.1016/j.cub.2009.08.044

26. Frichot, E., & François, O. (2015). LEA: An R package for landscape and ecological association studies. Methods in Ecology and Evolution, 6(8), 925–929. 10.1111/2041-210X.12382

27. Gallardo, B., Zieritz, A., Aldridge, D. C. (2015). The importance of the human footprint in shaping the global distribution of terrestrial, freshwater and marine invaders. PloS ONE, 10(5), e0125801. doi:10.1371/journal.pone.0125801

28. Gamba, C., Hanghøj, K., Gaunitz, C., Alfarhan, A. H., Alquraishi, S. A., Al-Rasheid, K. A. S., Bradley, D. G., & Orlando, L. (2016). Comparing the performance of three ancient DNA extraction methods for high-throughput sequencing. Molecular Ecology Resources, 16(2), 459–469. 10.1111/1755-0998.12470.

29. García-Verdugo, C., Caujapé-Castells, J., & Sanmartín, I. (2019). Colonization time on island settings: Lessons from the Hawaiian and Canary Island floras. Botanical Journal of the Linnean Society, 191(2), 155–163. 10.1093/botlinnean/boz044

30. Goodenough, A. (2010). Are the ecological impacts of alien species misrepresented? A review of the “native good, alien bad” philosophy. Community Ecology, 11(1), 13–21. 10.1556/ComEc.11.2010.1.3

31. Hume, J. P. (2005). Contrasting taphofacies in ocean island settings: the fossil record of Mascarene vertebrates. *In* Alcover, J.A. & Bover, P. (eds.): Proceedings of the International Symposium “Insular Vertebrate Evolution: the Palaeontological Approach”. Monografies de la Societat d’Història Natural de les Balears, 12, 129–144.

32. Hume, J. P. (2011). Systematics, morphology, and ecology of pigeons and doves (Aves: Columbidae) of the Mascarene Islands, with three new species. Zootaxa, 3124(1), 1. 10.11646/zootaxa.3124.1.1

33. Hume, J. P. (2013). A synopsis of the pre-human avifauna of the Mascarene Islands. In: Göhlich UB, Kroh A, editors. Proceed. 8th Internat. Meeting Society of Avian Paleontology and Evolution. Pp. 195–237. Wien, Naturhistorisches Museum.

34. Hume, J.P., de Louw, P.G.B., & Rijsdijk, K.F. (2015). Rediscovery of a lost Lagerstatte: a comparative analysis of the historical and recent Mare aux Songes dodo excavations on Mauritius. Historical Biology, 27(8), 1127–1140.

35. Johnson, K., Kort, S. D., Dinwoodey, K., Mateman, A. C., Cate, C. T., Lessells, C. M., & Clayton, D. H. (2001). A molecular phylogeny of the dove genera Streptopelia and Columba. 118(4), 874–887.

36. Johri, P., Aquadro, C. F., Beaumont, M., Charlesworth, B., Excoffier, L., Eyre-Walker, A., Keightley, P. D., Lynch, M., McVean, G., Payseur, B. A., Pfeifer, S. P., Stephan, W., & Jensen, J. D. (2021). Recommendations for improving statistical inference in population genomics. 10.1101/2021.10.27.466171

37. Jónsson, H., Ginolhac, A., Schubert, M., Johnson, P. L. F., & Orlando, L. (2013). mapDamage2.0: Fast approximate Bayesian estimates of ancient DNA damage parameters. Bioinformatics, 29(13), 1682–1684. 10.1093/bioinformatics/btt193

38. Kawakami, T., Smeds, L., Backström, N., Husby, A., Qvarnström, A., Mugal, C. F., Olason, P., & Ellegren, H. (2014). A high-density linkage map enables a second-generation collared flycatcher genome assembly and reveals the patterns of avian recombination rate variation and chromosomal evolution. Molecular Ecology, 23(16), 4035–4058. 10.1111/mec.12810

39. Kittelberger, K. D., Tanner, C. J., Buxton, A. N., Prewett, A., & Şekercioğlu, Ç. H. (2024). Correlates of avian extinction timing around the world since 1500 CE. Avian Research, 15, 100213. 10.1016/j.avrs.2024.100213

40. Korneliussen, T. S., Albrechtsen, A., & Nielsen, R. (2014). ANGSD: Analysis of Next Generation Sequencing Data. BMC Bioinformatics, 15(1), 356. 10.1186/s12859-014-0356-4

41. Leonardi, M., P. Librado, C. D. Sarkissian, M. Schubert, A. H. Alfarhan, S. A. Alquraishi, K. A. S. Al-Rasheid, C. Gamba, E. Willerslev and L. Orlando (2017). Evolutionary patterns and processes: lessons from ancient DNA. Systematic Biology 66(4): 660–660.

42. Lesturgie, P., Planes, S., & Mona, S. (2022). Coalescence times, life history traits and conservation concerns: An example from four coastal shark species from the Indo-Pacific. Molecular Ecology Resources, 22(2), 554–566. 10.1111/1755-0998.13487

43. Li, H., & Durbin, R. (2009). Fast and accurate short read alignment with Burrows–Wheeler transform. Bioinformatics, 25(14), 1754–1760. 10.1093/bioinformatics/btp324

44. Li, H., & Durbin, R. (2011). Inference of human population history from individual whole-genome sequences. Nature, 475(7357), 493–496. 10.1038/nature10231

45. Losos, J. B., & Ricklefs, R. E. (2009). Adaptation and diversification on islands. Nature, 457(7231), 830–836. 10.1038/nature07893

46. Marchi, N., Schlichta, F., & Excoffier, L. (2021). Demographic inference. Current Biology, 31(6), R276–R279. 10.1016/j.cub.2021.01.053

47. Martin, C. A., Armstrong, C., Illera, J. C., Emerson, B. C., Richardson, D. S., & Spurgin, L. G. (2021). Genomic variation, population history and within-archipelago adaptation between island bird populations. Royal Society Open Science, 8(2), rsos.201146, 201146. 10.1098/rsos.201146

48. Martin, C. A., Sheppard, E. C., Illera, J. C., Suh, A., Nadachowska-Brzyska, K., Spurgin, L. G., & Richardson, D. S. (2023). Runs of homozygosity reveal past bottlenecks and contemporary inbreeding across diverging populations of an island-colonizing bird. Molecular Ecology, 32(8), 1972–1989. 10.1111/mec.16865

49. Mather, N., Traves, S. M., & Ho, S. Y. W. (2020). A practical introduction to sequentially Markovian coalescent methods for estimating demographic history from genomic data. Ecology and Evolution, 10(1), 579–589. 10.1002/ece3.5888

50. Mazet, O., Rodríguez, W., Grusea, S., Boitard, S., & Chikhi, L. (2016). On the importance of being structured: Instantaneous coalescence rates and human evolution—lessons for ancestral population size inference? Heredity, 116(4), 362–371. 10.1038/hdy.2015.104

51. McKenna, A., Hanna, M., Banks, E., Sivachenko, A., Cibulskis, K., Kernytsky, A., Garimella, K., Altshuler, D., Gabriel, S., Daly, M., & DePristo, M. A. (2010). The Genome Analysis Toolkit: A MapReduce framework for analyzing next-generation DNA sequencing data. Genome Research, 20(9), 1297–1303. 10.1101/gr.107524.110

52. Meisner, J., & Albrechtsen, A. (2018). Inferring Population Structure and Admixture Proportions in Low-Depth NGS Data. Genetics, 210(2): 719–731.

53. Meyermans, R., Gorssen, W., Buys, N., & Janssens, S. (2020). How to study runs of homozygosity using PLINK? A guide for analyzing medium density SNP data in livestock and pet species. BMC Genomics, 21(1), 94. 10.1186/s12864-020-6463-x

54. Mills, L. S., & Allendorf, F. W. (1996). The One-Migrant-per-Generation Rule in Conservation and Management. Conservation Biology, 10(6), 1509–1518. 10.1046/j.1523-1739.1996.10061509.x

55. Mooney, H. A., & Cleland, E. E. (2001). The evolutionary impact of invasive species. Proceedings of the National Academy of Sciences, 98(10), 5446–5451. 10.1073/pnas.091093398

56. Myers, N., Mittermeier, R. A., Mittermeier, C. G., Da Fonseca, G. A. B., & Kent, J. (2000). Biodiversity hotspots for conservation priorities. Nature, 403(6772), 853–858. 10.1038/35002501

57. Nigenda-Morales, S. F., Lin, M., Nuñez-Valencia, P. G., Kyriazis, C. C., Beichman, A. C., Robinson, J. A., Ragsdale, A. P., Urbán R., J., Archer, F. I., Viloria-Gómora, L., Pérez-Álvarez, M. J., Poulin, E., Lohmueller, K. E., Moreno-Estrada, A., & Wayne, R. K. (2023). The genomic footprint of whaling and isolation in fin whale populations. Nature Communications, 14(1), 5465. 10.1038/s41467-023-40052-z

58. Palsbøll, P. J., Zachariah Peery, M., Olsen, M. T., Beissinger, S. R., & Bérubé, M. (2013). Inferring recent historic abundance from current genetic diversity. Molecular Ecology, 22(1), 22–40. 10.1111/mec.12094

59. Pemberton, T. J., Absher, D., Feldman, M. W., Myers, R. M., Rosenberg, N. A., & Li, J. Z. (2012). Genomic Patterns of Homozygosity in Worldwide Human Populations. The American Journal of Human Genetics, 91(2), 275–292. 10.1016/j.ajhg.2012.06.014

60. Privé, F., Luu, K., Vilhjálmsson, B. J., & Blum, M. G. B. (2020). Performing Highly Efficient Genome Scans for Local Adaptation with R Package pcadapt Version 4. Molecular Biology and Evolution, 37(7), 2153–2154. 10.1093/molbev/msaa053

61. Probst, J.-M. (2002). Fiche « patrimoine naturel à protéger » Le Ramier ou la Tourterelle malgache Streptopelia picturata. Revue d’Ecologie, 2002, 42–43.

62. Probst, J.-M., & Brial, P. (2002). Récits anciens de naturalistes à l’île Bourbon—Le 1er guide des espèces disparues de La Réunion (Reptiles, Oiseaux et Mammifères). Association Nature & patrimoine.

63. Quinlan, A. R., & Hall, I. M. (2010). BEDTools: A flexible suite of utilities for comparing genomic features. Bioinformatics, 26(6), 841–842. 10.1093/bioinformatics/btq033

64. Ragsdale, A. P., & Gravel, S. (2019). Models of archaic admixture and recent history from two-locus statistics. PLOS Genetics, 15(6), e1008204. 10.1371/journal.pgen.1008204

65. Reynolds, J., Weir, B. S., & Cockerham, C. C. (1983). Estimation of the Coancestry Coefficient: Basis for a Short-Term Genetic Distance. 105(3), 767–779. 10.1093/genetics/105.3.767

66. Rodriguez, L. F. (2006). Can Invasive Species Facilitate Native Species? Evidence of How, When, and Why These Impacts Occur. Biological Invasions, 8(4), 927–939. 10.1007/s10530-005-5103-3

67. Rodríguez, W., Mazet, O., Grusea, S., Arredondo, A., Corujo, J. M., Boitard, S., & Chikhi, L. (2018). The IICR and the non-stationary structured coalescent: Towards demographic inference with arbitrary changes in population structure. Heredity, 121(6), 663–678. 10.1038/s41437-018-0148-0

68. Roy, H. E., Pauchard, A., Stoett, P. J., Renard Truong, T., Meyerson, L. A., Bacher, S., Galil, B. S., Hulme, P. E., Ikeda, T., Kavileveettil, S., McGeoch, M. A., Nuñez, M. A., Ordonez, A., Rahlao, S. J., Schwindt, E., Seebens, H., Sheppard, A. W., Vandvik, V., Aleksanyan, A., … Ziller, S. R. (2024). Curbing the major and growing threats from invasive alien species is urgent and achievable. Nature Ecology & Evolution, 8(7), 1216–1223. 10.1038/s41559-024-02412-w

69. Russell, J. C., Meyer, J.-Y., Holmes, N. D., & Pagad, S. (2017). Invasive alien species on islands: Impacts, distribution, interactions and management. Environmental Conservation, 44(4), 359–370. 10.1017/S0376892917000297

70. Santiago, E., Novo, I., Pardiñas, A. F., Saura, M., Wang, J., & Caballero, A. (2020). Recent Demographic History Inferred by High-Resolution Analysis of Linkage Disequilibrium. Molecular Biology and Evolution, 37(12), 3642–3653. 10.1093/molbev/msaa169

71. Schlüter, T. (2016). Geological Atlas of Africa. Springer.

72. Schubert, M., Lindgreen, S., & Orlando, L. (2016). AdapterRemoval v2: Rapid adapter trimming, identification, and read merging. BMC Research Notes, 9(1), 88. 10.1186/s13104-016-1900-2

73. Seal, U. S., & Bruford, M. W. (1991). Columba (Nesoenas) Mayeri, Pink Pigeon. Captive Breeding Specialist Group (IUCN/SSC/CBSG).

74. Shapiro, M. D., Kronenberg, Z., Li, C., Domyan, E. T., Pan, H., Campbell, M., Tan, H., Huff, C. D., Hu, H., Vickrey, A. I., Nielsen, S. C. A., Stringham, S. A., Hu, H., Willerslev, E., Gilbert, M. T. P., Yandell, M., Zhang, G., & Wang, J. (2013). Genomic Diversity and Evolution of the Head Crest in the Rock Pigeon. Science, 339(6123), 1063–1067. 10.1126/science.1230422

75. Simberloff, D., Martin, J.-L., Genovesi, P., Maris, V., Wardle, D. A., Aronson, J., Courchamp, F., Galil, B., García-Berthou, E., Pascal, M., Pyšek, P., Sousa, R., Tabacchi, E., & Vilà, M. (2013). Impacts of biological invasions: What’s what and the way forward. Trends in Ecology & Evolution, 28(1), 58–66. 10.1016/j.tree.2012.07.013

76. Simberloff, D., Souza, L., Nuñez, M. A., Barrios-Garcia, M. N., & Bunn, W. (2012). The natives are restless, but not often and mostly when disturbed. Ecology, 93(3), 598–607. 10.1890/11-1232.1

77. Singhal, S., Leffler, E. M., Sannareddy, K., Turner, I., Venn, O., Hooper, D. M., Strand, A. I., Li, Q., Raney, B., Balakrishnan, C. N., Griffith, S. C., McVean, G., & Przeworski, M. (2015). Stable recombination hotspots in birds. 350(6263), 928–932. 10.1126/science.aad0843

78. Skoglund, P., Northoff, B. H., Shunkov, M. V., Derevianko, A. P., Pääbo, S., Krause, J., & Jakobsson, M. (2014). Separating endogenous ancient DNA from modern day contamination in a Siberian Neandertal. Proceedings of the National Academy of Sciences, 111(6), 2229–2234. 10.1073/pnas.1318934111

79. Spence, J. P., Steinrücken, M., Terhorst, J., & Song, Y. S. (2018). Inference of population history using coalescent HMMs: Review and outlook. Current Opinion in Genetics & Development, 53, 70–76. 10.1016/j.gde.2018.07.002

80. Spurgin, L. G., Illera, J. C., Jorgensen, T. H., Dawson, D. A., & Richardson, D. S. (2014). Genetic and phenotypic divergence in an island bird: Isolation by distance, by colonization or by adaptation? Molecular Ecology, 23(5), 1028–1039. 10.1111/mec.12672

81. Tajima, F. (1983). Evolutionary Relationship of DNA sequences in finite populations. Genetics, 105(2), 437–460. 10.1093/genetics/105.2.437

82. Tajima, F. (1989). Statistical Method for Testing the Neutral Mutation Hypothesis by DNA Polymorphism. 123(3), 585–595. 10.1093/genetics/123.3.585

83. Terhorst, J., Kamm, J. A., & Song, Y. S. (2017). Robust and scalable inference of population history from hundreds of unphased whole genomes. Nature Genetics, 49(2), 303–309. 10.1038/ng.3748

84. Thébaud, C., Strasberg, D., Warren, B. H., & Cheke, A. (2009). Mascarene Islands, Biology. In R. Gillespie & D. Clague (Eds.), Encyclopedia of Islands (pp. 612–619). University of California Press. 10.1525/9780520943728-146

85. Theissinger, K., Fernandes, C., Formenti, G., Bista, I., Berg, P. R., Bleidorn, C., Bombarely, A., Crottini, A., Gallo, G. R., Godoy, J. A., Jentoft, S., Malukiewicz, J., Mouton, A., Oomen, R. A., Paez, S., Palsbøll, P. J., Pampoulie, C., Ruiz-López, M. J., Secomandi, S., … Zammit, G. (2023). How genomics can help biodiversity conservation. Trends in Genetics, 39(7), 545–559. 10.1016/j.tig.2023.01.005

86. Van Der Auwera, G. A., Carneiro, M. O., Hartl, C., Poplin, R., Del Angel, G., Levy-Moonshine, A., Jordan, T., Shakir, K., Roazen, D., Thibault, J., Banks, E., Garimella, K. V., Altshuler, D., Gabriel, S., & DePristo, M. A. (2013). From FastQ Data to High-Confidence Variant Calls: The Genome Analysis Toolkit Best Practices Pipeline. Current Protocols in Bioinformatics, 43(1). 10.1002/0471250953.bi1110s43

87. Warren, B. H., Simberloff, D., Ricklefs, R. E., Aguilée, R., Condamine, F. L., Gravel, D., Morlon, H., Mouquet, N., Rosindell, J., Casquet, J., Conti, E., Cornuault, J., Fernández-Palacios, J. M., Hengl, T., Norder, S. J., Rijsdijk, K. F., Sanmartín, I., Strasberg, D., Triantis, K. A., … Thébaud, C. (2015). Islands as model systems in ecology and evolution: Prospects fifty years after MacArthur-Wilson. Ecology Letters, 18(2), 200–217. 10.1111/ele.12398

88. Watterson, W. (1975). On the number of segregating sites in genetical models without recombination. 7(2), 256–276. 10.1016/0040-5809(75)90020-9

89. Wittwer, S., Gerber, L., Allen, S. J., Willems, E. P., Marfurt, S. M., & Krützen, M. (2023). Reconstructing the colonization history of INDO-PACIFIC bottlenose dolphins ( *Tursiops aduncus* ) in Northwestern Australia. Molecular Ecology, 32(14), 3826– 3841. 10.1111/mec.16984

90. Wolfenden, A., Jones, C. G., Tatayah, V., Züel, N., & De Kort, S. R. (2015). Endangered pink pigeons treat calls of the ubiquitous Madagascan turtle dove as conspecific. Animal Behaviour, 99, 83–88. 10.1016/j.anbehav.2014.10.023

91. Zhang, G., Li, C., Li, Q., Li, B., Larkin, D. M., Lee, C., Storz, J. F., Antunes, A., Greenwold, M. J., Meredith, R. W., Ödeen, A., Cui, J., Zhou, Q., Xu, L., Pan, H., Wang, Z., Jin, L., Zhang, P., Hu, H., … Froman, D. P. (2014). Comparative genomics reveals insights into avian genome evolution and adaptation. Science, 346(6215), 1311–1320. 10.1126/science.1251385

