## Supplementary Material for "Combining population genomics with ancient DNA to understand island colonization history of the Madagascar turtle dove"

### S1: Reference genome sequencing and assembly

Long-read PacBio sequencing was performed on two SMRTcells at the ICGex platform of the Curie institute (Paris). We generated approximately 90 million reads, equalling an estimated ~90X coverage. After converting the BAM sequencing files to FASTQ using bedtools, we performed genome assembly using Hifiasm (version 0.16.1-r375). Total assembly size was 1.25 Gb across 409 contigs, with a contig N50 of 33.1 Mb. We excluded potential sex chromosomes from downstream analyses using the nucmer program in MUMmer (v.4.0.0beta2; Marçais et al., 2018) to perform pairwise alignments between each scaffold and the W and Z chromosome of the domestic rock pigeon (*Columba livia*), the closest species with a fully annotated genome (NCBI RefSeq: GCF_036013475.1). We ran RepeatMasker v4.1.0 to scan the genome for repetitive elements present in the *de novo* and *Aves* database (Smit et al., 2013). RepeatModeler v2.0.1 was further used to create a *de novo* database of repetitive elements specific to *N. picturata* (Smit & Hubley, 2008). Repetitive elements in the genome were conserved in a BED file and excluded from downstream analyses.

### S2: Ancient DNA methods

*DNA extraction and sample library preparation*

DNA extraction followed Gamba et al. (2016) with modifications from Fages et al. (2019). In detail, the standard protocol makes use of 1mL of lysis buffer (0.45M EDTA, 0.25 mg/mL proteinase K and 0.5% sodium lauroyl sarcosinate) to carry out a first pre-digestion step of 30 min at 37°C. Following centrifugation at 3,000 rpm for 2 min, the supernatant was removed and a 1 mL volume of a fresh lysis buffer was added, followed by full digestion of the remaining pellets at 42°C overnight. After centrifugation at 3,800 rpm for 2 min, the supernatant was concentrated using Amicon Ultra-4 columns (Millipore) and 250 µmL of the remaining supernatant was purified through MinElute columns (QIAgen) at a 1:10 ratio (supernatant:PB buffer) following manufacturer's instructions, and concentrated in a 60 µL of elution buffer (Qiagen EB Buffer supplemented with 0.05 % Tween 20).

Two different sets of libraries were produced per sample. One library per sample was built from non-USER-treated ancient DNA, amplified with one PCR reaction for low coverage shotgun sequencing on an inhouse Illumina MiniSeq instrument (using 2 × 80 bp High-Output Kit) to assess the damage of ancient DNA and endogenous DNA content (Seguin-Orlando et al. 2021). Based on these initial shotgun sequencing analyses, if endogenous DNA in the sample was deemed to be of sufficient quality and quantity, a second library per sample was built from the USER-treated ancient DNA by adding 7 µL of USER enzyme (NEB) to a total of 22.8 µL purified extract for 3 hours at 37°C in order to remove post mortem DNA damage (Gaunitz et al., 2018). This second library was then amplified specifically for hyRAD capture and deep-sequencing.

A total of 14.9 µL of ancient DNA extract (USER-treated or not) was used for library preparation following a modified version of the Meyer & Kircher (2010) protocol presented in Fages et al. (2019)**,** including one 6-bp “external” index (Orlando et al., 2013) and two 7-bp “internal” indices (Rohland et al., 2015). Before hyRAD capture, each of the USER-treated ancient DNA libraries prepared were amplified in two successive rounds of eight parallel PCRs (Clavel et al.,2023). These rounds ensured sufficient amplification necessary for hyRAD capture (Suchan et al. 2016, 2022) without compromising library diversity. Depending on the initial PCR round, the number of PCR cycles and parallel runs for the second round were increased or reduced.

The first round was amplified in eight parallel PCRs for eight cycles at the same conditions: a total reaction volume of 25 µL, using 0.4 µL of AccuPrime Pfx DNA polymerase (one unit), 2.5 µL 10X Accuprime Pfx reaction mix, 2 µL of DNA library, 1 µL of BSA (20 mg/mL), 17.9 µL of H2O, 0.2 µL of inPE1 primer (25 mM) and 1 µL index primer (5 mM) containing a unique 6-bp “external” barcode (as explained above). Two amplification products for a given library were then merged and purified together using Agencourt Ampure XP beads (1:1.2 or 1:1.4 as DNA:beads ratio), and eluted in 11 µL of EB-Tween buffer. This provided four purified amplification products per library, which were then split into two parallel PCR amplifications, using 1 µL of the eluate for each PCR reaction, for 10 cycles. The second PCR round was carried out in 25 µL reaction volumes, following the same conditions as for the first amplification round, except that IS5 and IS6 were used as PCR primers (see Meyer et Kircher 2010). The resulting eight PCR products were co-purified in a single MinElute column (QIAGEN), and eluted in 15 µL of EB-Tween buffer. Library concentration and size distribution were measured throughout the process on a Qubit HS dsDNA assay (Invitrogen) and on a Tapestation 4200 instrument using a High sensitivity D1000 ScreenTapeAssay (Agilent technologies).

*HyRAD probe library preparation and probe production*

Probe production was based on the previous hyRAD protocol (Suchan et al., 2016, 2022; full protocol available: dx.doi.org/10.17504/protocols.io.bywxpxfn), which is based on the development of a RADseq library and relies on the presence of a T7 RNA polymerase promoter in the DNA probe library adapter to allow *in vitro* transcription to RNA probes. Various restriction enzymes were tested through simulated digestion of the *Neosenas mayeri* genome with the aim of including approximately 10% of the genome within the probe set. The *PstI-MseI* restriction enzyme combination which targets 6bp- and 4bp-long restriction sites, respectively, was identified to be most relevant, since it was expected to produce around 500,000 digested fragments within the 100-300 bp probe design range.

Fresh high-molecular weight DNA extracts (500 to 1,000 ng) from six *Nesoenas mayeri* and six *Nesoenas picturata* blood samples were subject to *PstI-MseI* double enzymatic restriction before being ligated to two adapters, each showing terminal ends complementary to one restriction site. Each digestion reaction (80 µl) included 40 U of *PstI*-HF (NEB) and 20 U of *MseI* (NEB) in the CutSmart buffer for 3h at 37°C. The reaction was purified using AMPure beads (Beckman Coulter), with a bead-to-liquid ratio of 2:1 and eluted in 20 µl of 10 mM Tris-HCL. The purified DNA was then adapter ligated through a 3h ligation reaction at 37°C with 800 U of T4 DNA ligase (NEB) and 0.7 µM of each adapter (RAD-P1-T7-PstI and RAD-P2-MseI, with a final volume of 28 µl. Following purification with AMPure beads (bead-to-liquid ratio = 1.5:1) and elution in 60 µL of 10 mM Tris-HCL, DNA templates were selected within the 180-380 bp size range using the BluePippin instrument (Sage Science) and 2% agarose cassettes with external marker (Sage Science).  Given that the total size of the adapters represents 90 bp, the selected size range corresponds to inserts between approximately 90-290 bp, as per Suchan et al., (2022). Size ranges were monitored using the TapeStation 4200 (Agilent) prior and post size selection. A follow-up purification step (biotinylated fragment selection) allowed the selection of DNA fragments containing the aforementioned adapters (specifically targeting the RAD-P2 adapter containing the biotin molecule) using streptavidin-coated beads (MyOne C1 Dynabeads, Invitrogen). A total of 30 µL of the beads was first washed and resuspended in 40µL of 2x Bind and Wash buffer, then combined with 40 µL of DNA and  incubated whilst rotating for 15 min. This was subsequently separated on a magnetic rack, washed three times with 1x Bind and Wash buffer, and resuspended in 30 µL of 10mM Tris-HCL with the beads.

The resultant probe library constructs were PCR amplified in a single reaction using KAPA HiFi HotStart ReadyMix (Roche) and 0.6 µM of the IS4 primer and indexing primers per specimen (Meyer & Kircher, 2010). The PCR conditions were an initial denaturation of 3 min at 95°C, 15 cycles of 20 s at 98°c, 15 s at 60°C, 30 s at 72°C, and final extension of 5 min at 72°C. The probe library was purified using AMPure beads (bead-to-liquid ratio = 1:1), eluted in 20 µL of 10 mM Tris-HCL and quantified using Qubit. Adapter removal was performed by digesting the purified probe library with 1.5 U of *MseI* in a 15 µL reaction for 3 h at 37°C. Adapter removal was confirmed by running the sample on TapeStation 4200 instrument (Agilent) and comparing results prior to and after digestion. Adapter removed purified probe libraries of 11 out of the 12 *Neosenas* samples (one was removed due to low concentration) were subsequently pooled in equimolar portions.

In a final step, RNA probes were synthesized through an *in vitro* transcription reaction using HiScribe T7 High Yield RNA Synthesis Kit (NEB) according to the manufacturer's instructions. This included incubating biotin-16-UTP (1/3 of UTP molarity; Roche), other nucleotides (GTP, ATP, CTP), T7 RNA polymerase, and 6.5 µl of the equimolar pooled probe library DNA template in a 20 µl reaction for 16 h at 37°C. Each reaction was then subject to TurboDNAse (4U; Thermo Fisher) treatment at 37°C for 30 min in order to remove remaining DNA templates and then purified on RNEasy Mini column (Qiagen) using the standard procedure, with the addition of 665 µl of Ethanol to the RNA and RTL buffer mix before loading it into the column. Final elution was made in 25 µl of RNase-free water and RNA probes were diluted to 100 ng/µL. Finally, for every 19 uL of RNA probes, 1 uL of SUPERase-In RNase inhibitor (20U; Thermo Fisher) was added. HyRAD capture requires the use of RNA blocking oligonucleotides (from herein, blockers) that are complementary to the Illumina P5 and P7 adapters present in sample DNA libraries. In order to synthesize such blockers, we annealed synthetic BO.P5 and BO.P7 DNA oligonucleotides with a synthetic oligonucleotide consisting of the P7 promoter sequence (See Table S1 in Suchan et al 2022; Carpenter et al., 2013) at 100 µM concentration. We then used 1 µL of each annealed template (P5 and P7) in separate transcription reactions using the same HiScribe T7 High Yield RNA Synthesis Kit (NEB) process above, includingTurboDNAse treatment, RNEasy Mini column purification, and the addition of SUPERase-In.

*Hybridization capture*

Hybridization conditions closely followed protocols of MyBaits v3 (MYcroarray, USA; http://www.mycroarray.com/mybaits/manuals.html) and Suchan et al. (2016 and 2022). However, we used 1000 ng of input DNA instead of 500 ng. For each reaction, each USER-treated DNA library was mixed with a blocking mix consisting of 0.55 µM of each blocker (P7 and P5), 2.3 µg of human Cot-1 DNA, and 2.3 µg of salmon sperm DNA (from herein, LYBs mix). The hybridization reaction mix included: 5.4x SSPE (equivalent to 0.8 M NaCl), 0.15% SDS, 5.25x Denhardt's solution, 0.9 U of SUPERase-In (Thermo Fisher), 13 mM EDTA (including the one present in the SSPE buffer), and 500 ng of RNA probes (from herein, HYBs mix). The LYBs mix was denatured for 5 minutes at 95°C, then the temperature was lowered to 55°C at which the prewarmed HYBs mix (at 55°C) was added to the LYBs mix to hybridize for 24 hours (Cruz-Dávalos et al., 2017).

Hybridized DNA fragments were captured during a 30-minute incubation at 55°C with 30 µl of streptavidine-coated beads (Dynabeads C1, Thermo Fisher) resuspended in 70 µl of TEN buffer (10 mM Tris-HCl pH 7.5, 1 mM EDTA, 1M NaCl). Next, beads containing the hybridized DNA fragments were separated using a magnetic rack, washed through resuspension and incubation cycles for 15 minutes in 180 µl of 1x SSC/0.1% SDS (once) and for 10 minutes with 180 µl of 0.1x SSC/0.1% SDS (three times), before being resuspended in 30 µl of molecular grade water. If required, we carried out a second hybridization capture, of 16h to 24h following the same protocol above.

Amplification reactions were performed in duplicates (two PCR runs per sample) using KAPA HiFi HotStart ReadyMix, 7.5 µL of the beads solution after the capture and wash steps, and 0.5 µM of IS5_reamp.P5 and IS6_reamp.P7 primers (see Suchan et al., 2022) from Meyer & Kircher (2010). These amplification reactions were either between rounds of capture (12 PCR cycles- aimed at 500 ng of input DNA for the second round or amplification for sequencing (8 PCR cycles - whether first or second round of capture). All resulting captured samples were sequenced at low coverage on an inhouse Illumina MiniSeq instrument using a 2 × 80 bp High-Output Kit for initial analysis of capture success and endogenous content. Deep coverage sequencing was subsequently performed using the Illumina NextSeq instrument using a 2 x 100bp or 2 x 150 bp run (iGenSeq platform, ICM Paris).

### S3: Obtaining the IICR and computation of LD decay

The simulated *IICR(t)* is obtained from the following equation:

$$IICR(t_i) = \frac{1 - F_{T_2(t_i)}}{f_{T_2}(t_i)} \quad \text{(Rodríguez et al., 2018)}$$

We simulated expected LD under our most likely model. LD measures the correlation between alleles at different variant sites and decreases over time with recombination. The recombination rate depends on the physical distance between variants and the effective population size. Consequently, LD can be used to infer demographic history (Fournier et al., 2023; Ragsdale & Gravel, 2019; Santiago et al., 2020). The *moments.LD* program computes the expectation of LD decay under complex multi-population scenarios (Ragsdale & Gravel, 2019, 2020). Specifically, it computes $\mathbb{E}[D^2]$ and $\mathbb{E}[D_z]$ for each deme in a given model. $\mathbb{E}[D^2]$ represents the variance of the two-locus statistic D, a measure of LD that quantifies the covariance of allele frequencies between two loci (Ragsdale & Gravel, 2019). $\mathbb{E}[D_z]$ is the expectation of positive covariance among low-frequency variants (Ragsdale & Gravel, 2019). Both LD statistics are normalized by $\mathbb{E}[\pi_2]$, the expectation of joint heterozygosity across SNP pairs (Ragsdale & Gravel, 2019).

### References S1-S3


Carpenter, M. L., Buenrostro, J. D., Valdiosera, C., Schroeder, H., Allentoft, M. E., Sikora, M., Rasmussen, M., Gravel, S., Guillén, S., Nekhrizov, G., Leshtakov, K., Dimitrova, D., Theodossiev, N., Pettener, D., Luiselli, D., Sandoval, K., Moreno-Estrada, A., Li, Y., Wang, J., … Bustamante, C. D. (2013). Pulling out the 1%: Whole-Genome Capture for the Targeted Enrichment of Ancient DNA Sequencing Libraries. *The American Journal of Human Genetics*, *93*(5), 852–864. https://doi.org/10.1016/j.ajhg.2013.10.002.

Clavel P, Louis L, Sarkissian C, Thèves C, Gillet C, Chauvey L, Tressières G, Schiavinato S, Calvière-Tonasso L, Telmon N, Clavel B, Jonvel R, Tzortzis S, Bouniol L, Fémolant JM, Klunk J, Poinar H, Signoli M, Costedoat C, Spyrou MA, Seguin-Orlando A, Orlando L. (2023). Improving the extraction of ancient *Yersinia pestis* genomes from the dental pulp. *iScience*, 2;26(5):106787. doi: 10.1016/j.isci.2023.106787.

Cruz-Dávalos, D. I., Llamas, B., Gaunitz, C., Fages, A., Gamba, C., Soubrier, J., Librado, P., Seguin-Orlando, A., Pruvost, M., Alfarhan, A. H., Alquraishi, S. A., Al-Rasheid, K. A. S., Scheu, A., Beneke, N., Ludwig, A., Cooper, A., Willerslev, E., & Orlando, L. (2017). Experimental conditions improving in-solution target enrichment for ancient DNA. *Molecular Ecology Resources*, *17*(3), 508–522. https://doi.org/10.1111/1755-0998.12595.

Fages, A., Hanghøj, K., Khan, N., Gaunitz, C., Seguin-Orlando, A., Leonardi, M., McCrory Constantz, C., Gamba, C., Al-Rasheid, K. A. S., Albizuri, S., Alfarhan, A. H., Allentoft, M., Alquraishi, S., Anthony, D., Baimukhanov, N., Barrett, J. H., Bayarsaikhan, J., Benecke, N., Bernáldez-Sánchez, E., … Orlando, L. (2019). Tracking Five Millennia of Horse Management with Extensive Ancient Genome Time Series. *Cell*, *177*(6), 1419-1435.e31. https://doi.org/10.1016/j.cell.2019.03.049.

Fournier, R., Tsangalidou, Z., Reich, D., & Palamara, P. F. (2023). Haplotype-based inference of recent effective population size in modern and ancient DNA samples. *Nature Communications*, *14*(1), 7945. https://doi.org/10.1038/s41467-023-43522-6

Gamba, C., Hanghøj, K., Gaunitz, C., Alfarhan, A. H., Alquraishi, S. A., Al-Rasheid, K. A. S., Bradley, D. G., & Orlando, L. (2016). Comparing the performance of three ancient DNA extraction methods for high-throughput sequencing. *Molecular Ecology Resources*, *16*(2), 459–469. https://doi.org/10.1111/1755-0998.12470.

Gaunitz, C., Fages, A., Hanghøj, K., Albrechtsen, A., Khan, N., Schubert, M., Seguin-Orlando, A., Owens, I.J., Felkel, S., Bignon-Lau, O., et al. (2018). Ancient genomes revisit the ancestry of domestic and Przewalski's horses. Science 360, 111–114. https://doi.org/10.1126/science. aao3297.

Marçais, G., Delcher, A. L., Phillippy, A. M., Coston, R., Salzberg, S. L., & Zimin, A. (2018). MUMmer4: A fast and versatile genome alignment system. *PLOS Computational Biology*, *14*(1), e1005944. https://doi.org/10.1371/journal.pcbi.1005944

Meyer, M., & Kircher, M. (2010). Illumina Sequencing Library Preparation for Highly Multiplexed Target Capture and Sequencing. *Cold Spring Harbor Protocols*, *2010*(6), pdb.prot5448. https://doi.org/10.1101/pdb.prot5448.

Orlando, L., Ginolhac, A., Zhang, G., Froese, D., Albrechtsen, A., Stiller, M., Schubert, M., Cappellini, E., Petersen, B., Moltke, I., Johnson, P. L. F., Fumagalli, M., Vilstrup, J. T., Raghavan, M., Korneliussen, T., Malaspinas, A.-S., Vogt, J., Szklarczyk, D., Kelstrup, C. D., … Willerslev, E. (2013). Recalibrating Equus evolution using the genome sequence of an early Middle Pleistocene horse. *Nature*, *499*(7456), 74–78. https://doi.org/10.1038/nature12323.

Ragsdale, A. P., & Gravel, S. (2019). Models of archaic admixture and recent history from two-locus statistics. *PLOS Genetics*, *15*(6), e1008204. https://doi.org/10.1371/journal.pgen.1008204

Ragsdale, A. P., & Gravel, S. (2020). Unbiased Estimation of Linkage Disequilibrium from Unphased Data. *Molecular Biology and Evolution*, *37*(3), 923–932. https://doi.org/10.1093/molbev/msz265

Rohland, N., Harney, E., Mallick, S., Nordenfelt, S., & Reich, D. (2015). Partial uracil–DNA–glycosylase treatment for screening of ancient DNA. *Philosophical Transactions of the Royal Society B: Biological Sciences*, *370*(1660), 20130624. https://doi.org/10.1098/rstb.2013.0624.

Santiago, E., Novo, I., Pardiñas, A. F., Saura, M., Wang, J., & Caballero, A. (2020). Recent Demographic History Inferred by High-Resolution Analysis of Linkage Disequilibrium. *Molecular Biology and Evolution*, *37*(12), 3642–3653. https://doi.org/10.1093/molbev/msaa169

Seguin-Orlando, A., Costedoat, C., Der Sarkissian, C., Tzortzis, S., Kamel, C., Telmon, N., Dale ́ n, L., The ̀ ves, C., Signoli, M., and Orlando, L. (2021). No particular genomic features underpin the dramatic economic consequences of 17th century plague epidemics in Italy. iScience 24, 102383. https://doi.org/10.1016/j.isci.2021.
102383.

Schmid, S., Genevest, R., Gobet, E., Suchan, T., Sperisen, C., Tinner, W., & Alvarez, N. (2017). HyRAD-X, a versatile method combining exome capture and RAD sequencing to extract genomic information from ancient DNA. *Methods in Ecology and Evolution*, *8*(10), 1374–1388. https://doi.org/10.1111/2041-210X.12785.

Smit, A., & Hubley, R. (2008). *RepeatModeler Open-1.0* [Computer software]. http://www.repeatmasker.org

Smit, A., Hubley, R., & Green, P. (2013). *RepeatMasker Open-4.0* [Computer software]. http://www.repeatmasker.org

Suchan, T., Pitteloud, C., Gerasimova, N. S., Kostikova, A., Schmid, S., Arrigo, N., Pajkovic, M., Ronikier, M., & Alvarez, N. (2016). Hybridization Capture Using RAD Probes (hyRAD), a New Tool for Performing Genomic Analyses on Collection Specimens. *PLOS ONE*, *11*(3), e0151651. https://doi.org/10.1371/journal.pone.0151651.

Suchan T, Kusliy MA, Khan N, Chauvey L, Tonasso-Calvière L, Schiavinato S, Southon J, Keller M, Kitagawa K, Krause J, Bessudnov AN, Bessudnov AA, Graphodatsky AS, Valenzuela-Lamas S, Wilczyński J, Pospuła S, Tunia K, Nowak M, Moskal-delHoyo M, Tishkin AA, Pryor AJE, Outram AK, Orlando L. (2022) Performance and automation of ancient DNA capture with RNA hyRAD probes. Mol Ecol Resour.;22(3):891-907. doi: 10.1111/1755-0998.13518.

**Figure S1:** Representation of the eight models evaluated in *fastsimcoal2*, grouped by the hypothesis being tested. **(A)** Testing for tree topology and colonization sequence (MOD1→MOD6). We limited model scope with geography and the results of population structure making Madagascar the ancestral deme in all models except MOD1, a scenario that could cause the observed sNMF clustering (see Main Text). **(B)** Testing for gene flow. We incorporate gene flow (as asymmetric migrations) to the most likely tree topology. (**C)** Testing for changes in *Ne* and recent translocations from Madagascar. This model incorporates information obtained from historical records and demographic inferences.

**A)** Testing for tree topology and colonization sequence

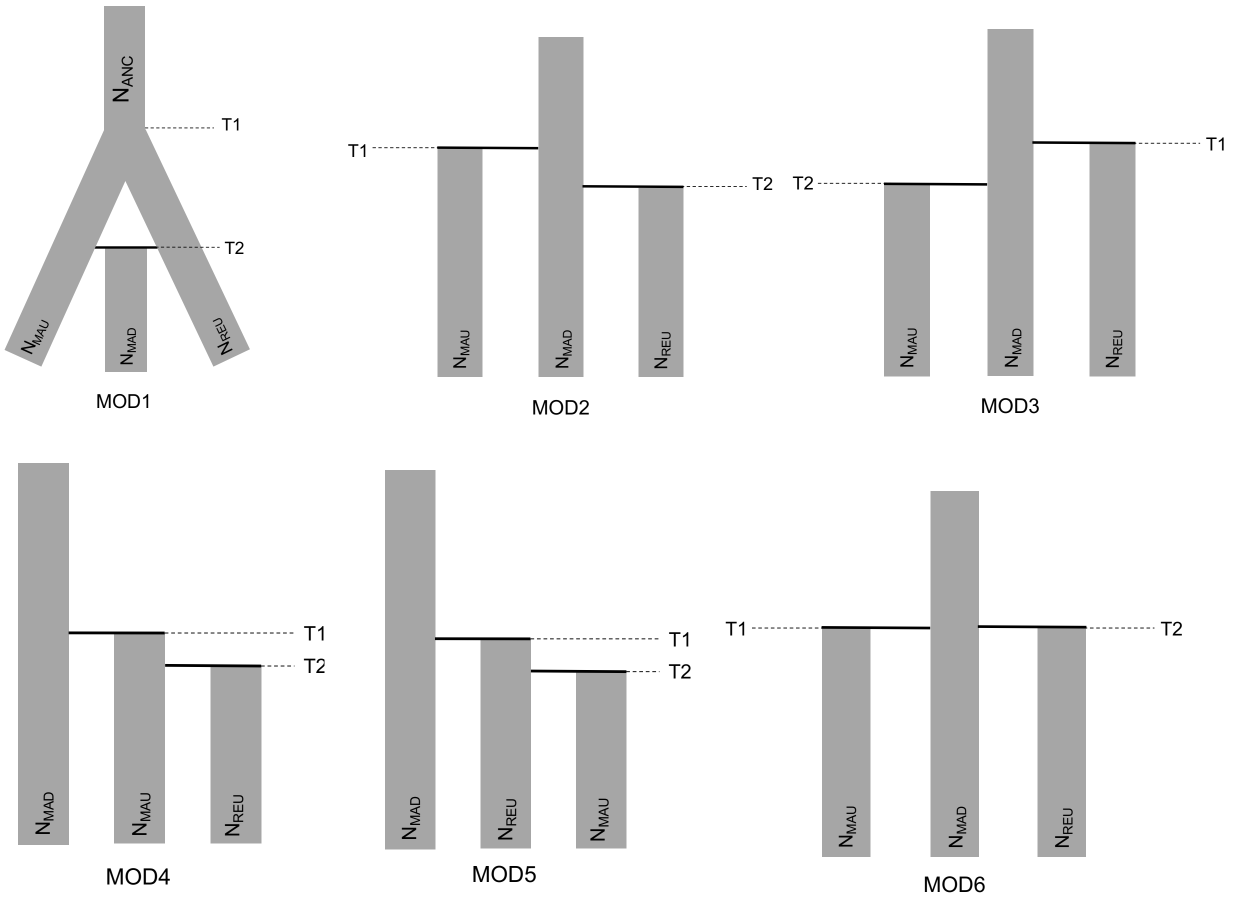

**B**) Testing for gene flow

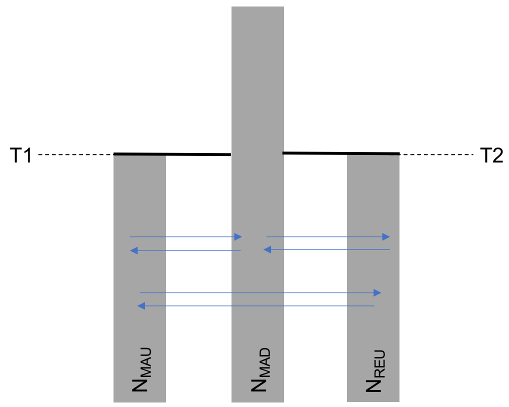


MOD6_MIG

**C)** Testing for changes in *Ne* and recent translocations from Madagascar

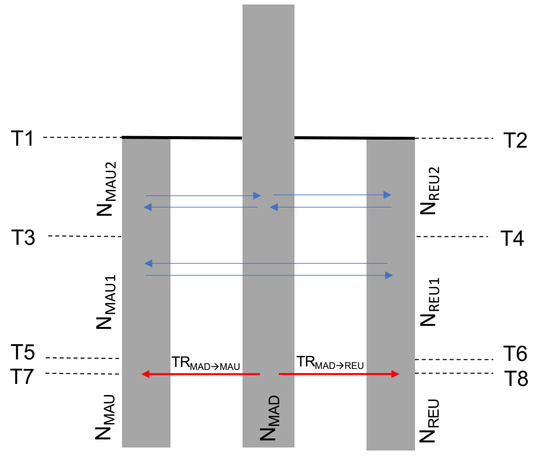


MOD6_MIG_TR

**Figure S2:** Heatmap representing p-distances between *N. picturata* individuals.

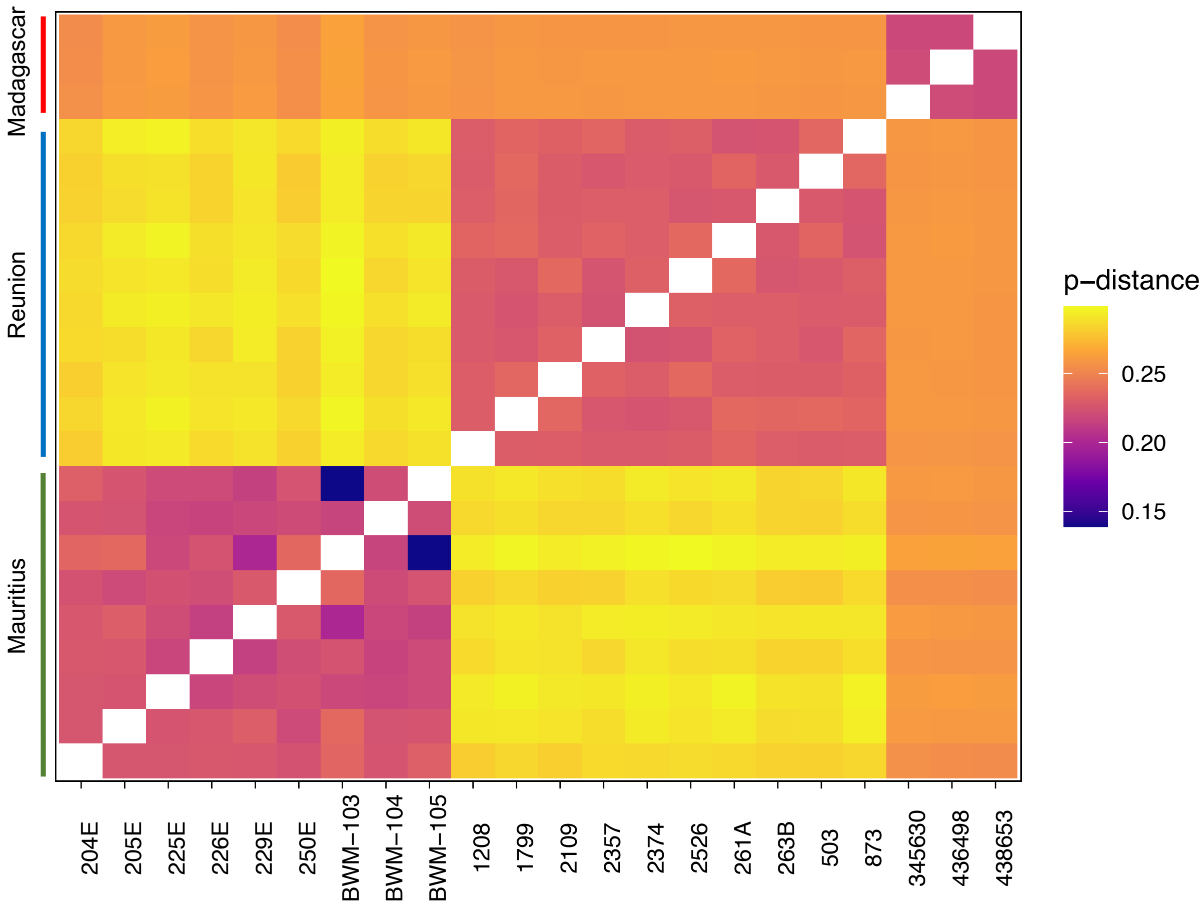

**Figure S3:** Cross-validation of the sNMF clustering algorithm on the *N. picturata* dataset for ancestral populations ranging from K=1 to K=7.

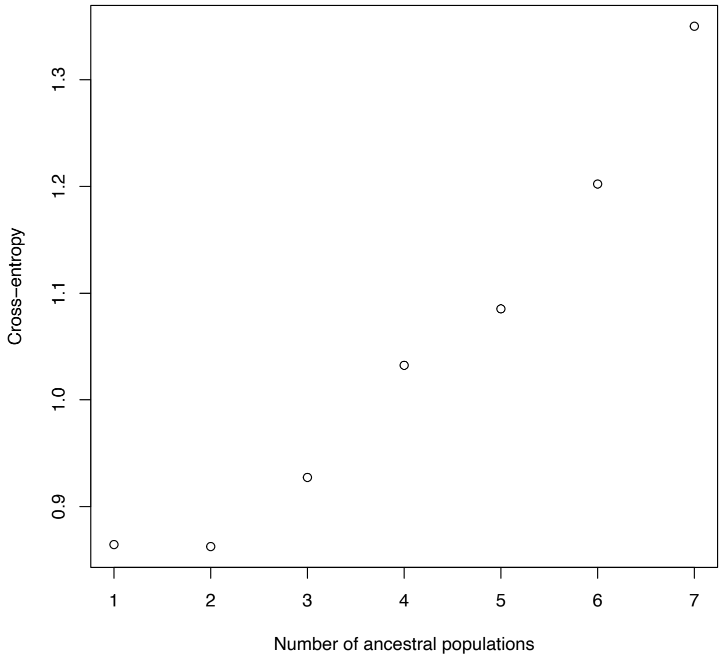

**Figure S4:** Heatmap showing the pairwise IBS distance between the shared sequences of subfossil Sub1, morphologically identified as likely *N. cicur* or *N. picturata*, and modern *N. picturata*.

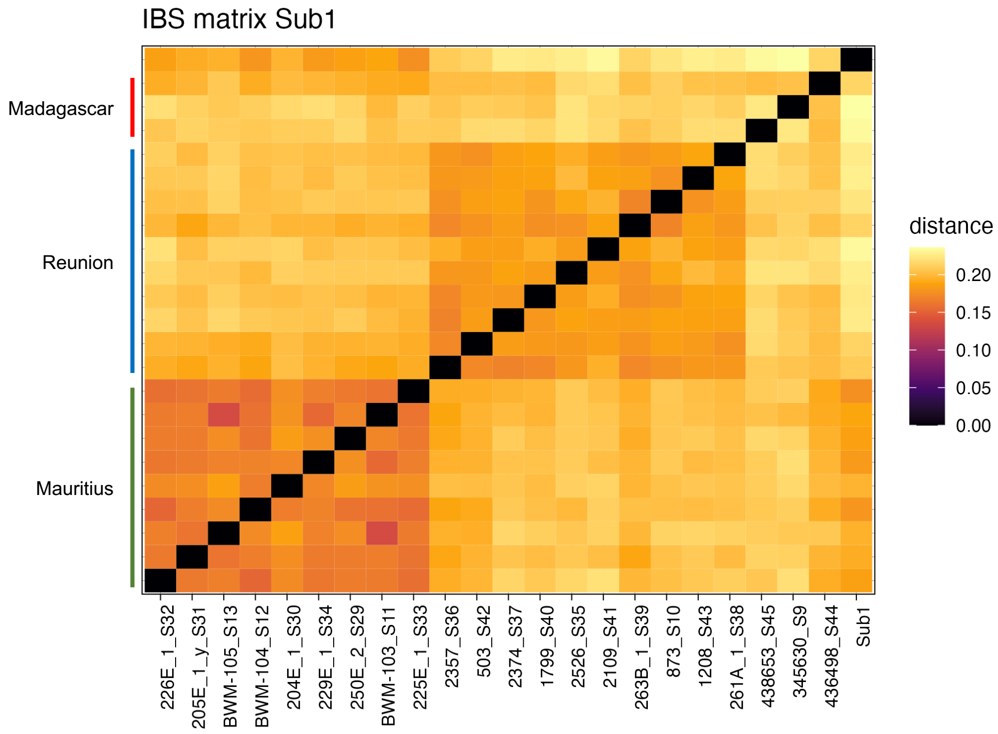

**Figure S5:** Heatmap showing the pairwise IBS distance between the shared sequences of subfossil Sub13, morphologically identified as likely *N. picturata*, and modern *N. picturata*.

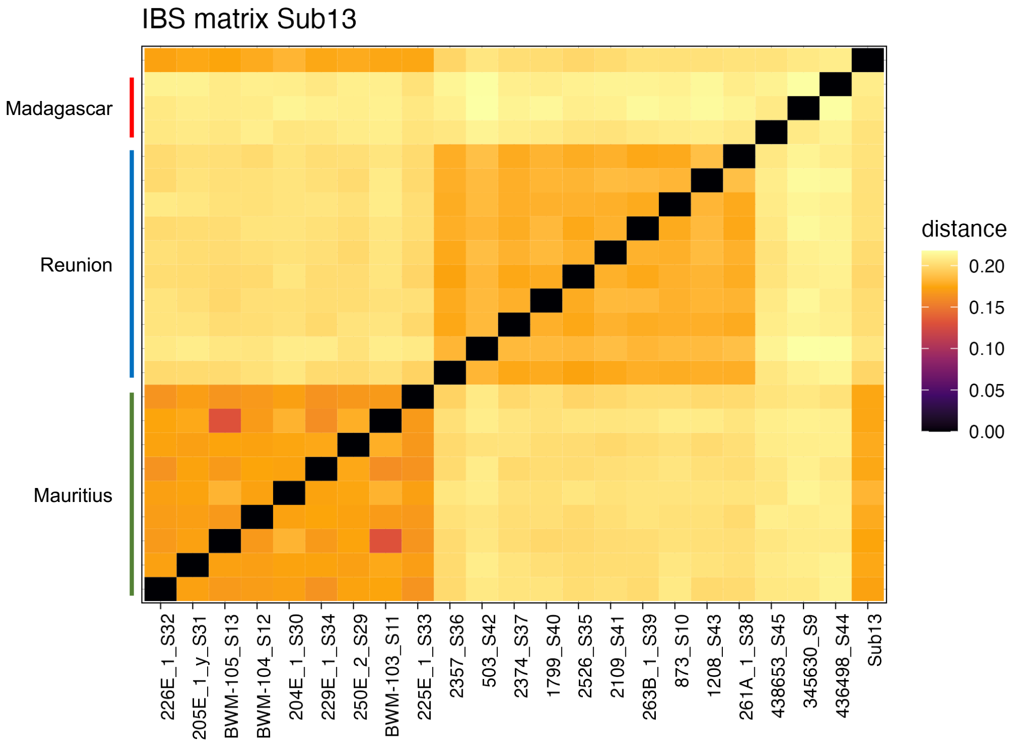

**Figure S6:** Heatmap showing the pairwise IBS distance between the shared sequences of subfossil Sub4_S11, morphologically identified as *N. cicur* or *N. picturata*, and modern *N. picturata*.

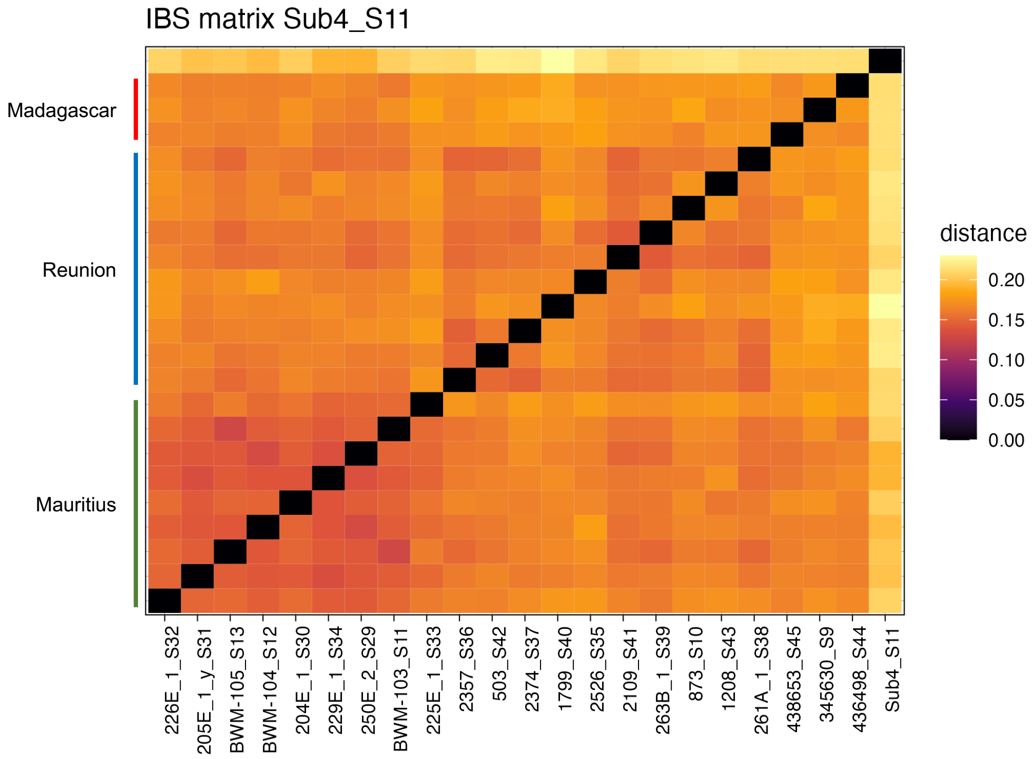

**Figure S7:** Heatmap showing the pairwise IBS distance between the shared sequences of subfossil Sub3, morphologically identified as *N. cicur* or *N. picturata,* and modern *N. picturata*.

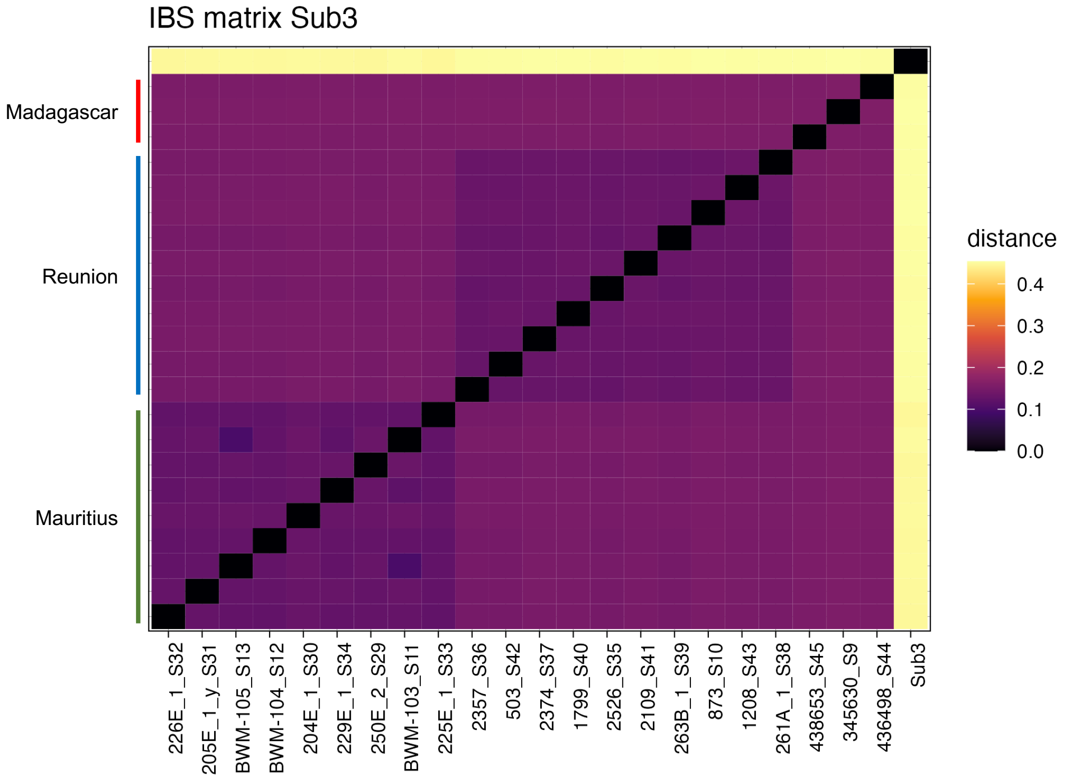

**Figure S8**: Genetic diversity in *N. picturata* populations measured by the average number of pairwise differences between sequences, $\theta_\pi$. Boxes represent the distribution over the 32 scaffolds longer than 10M bp.

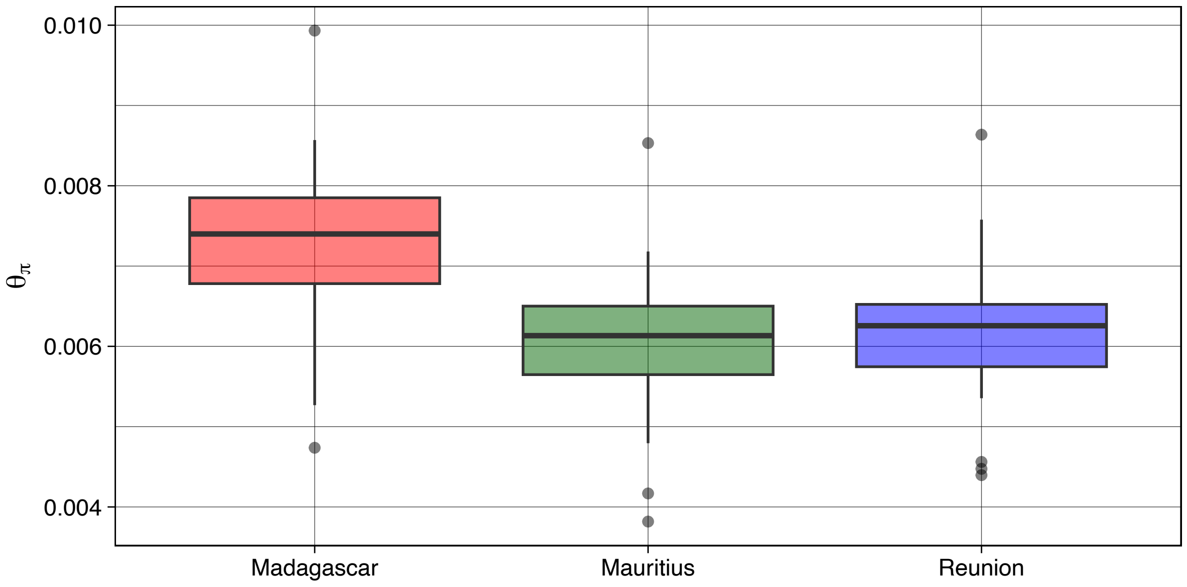

**Figure S9:** Number of heterozygous positions in 300k bp windows of scaffold 1 (10,934,710 bp) per *N. picturata* individual. We include one individual per population as examples in addition to the anomalous Mauritian individuals, 205E and 225E. Colours represent the populations of origin; red= Madagascar, blue=Reunion, green=Mauritius.

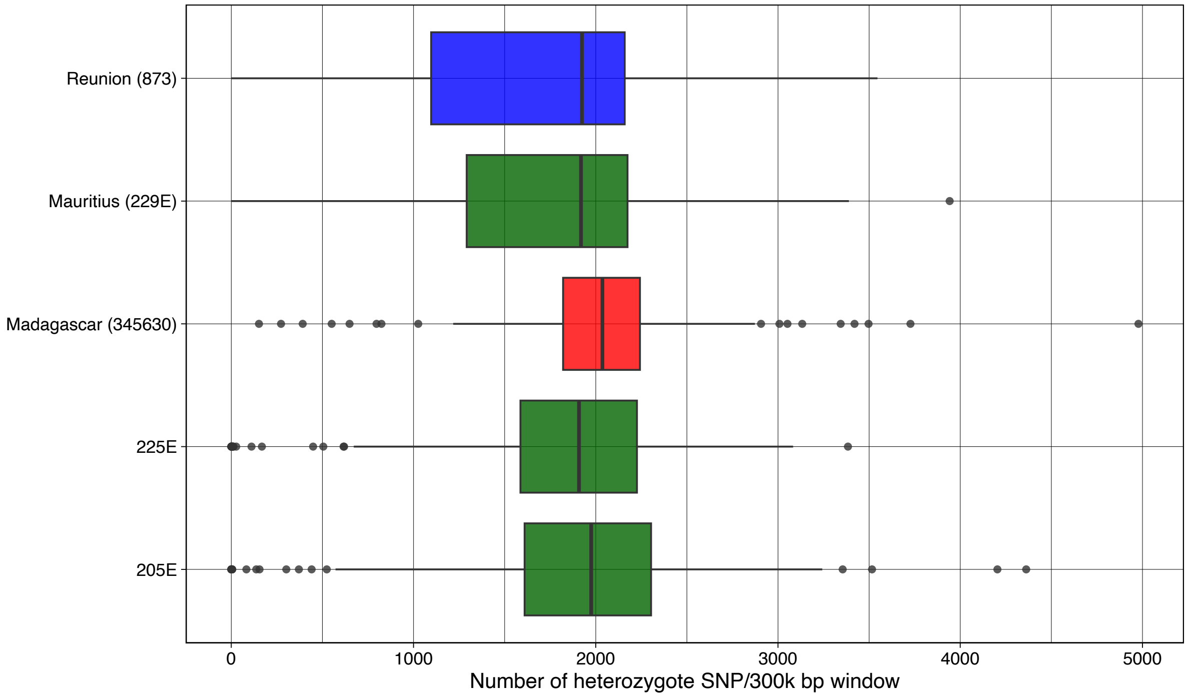

**Figure S10:** Intra-population PCA. Each point represents an individual labelled by its samplename. The populations are **(A)** Mauritius (N=9). **(B)** Reunion (N=10).

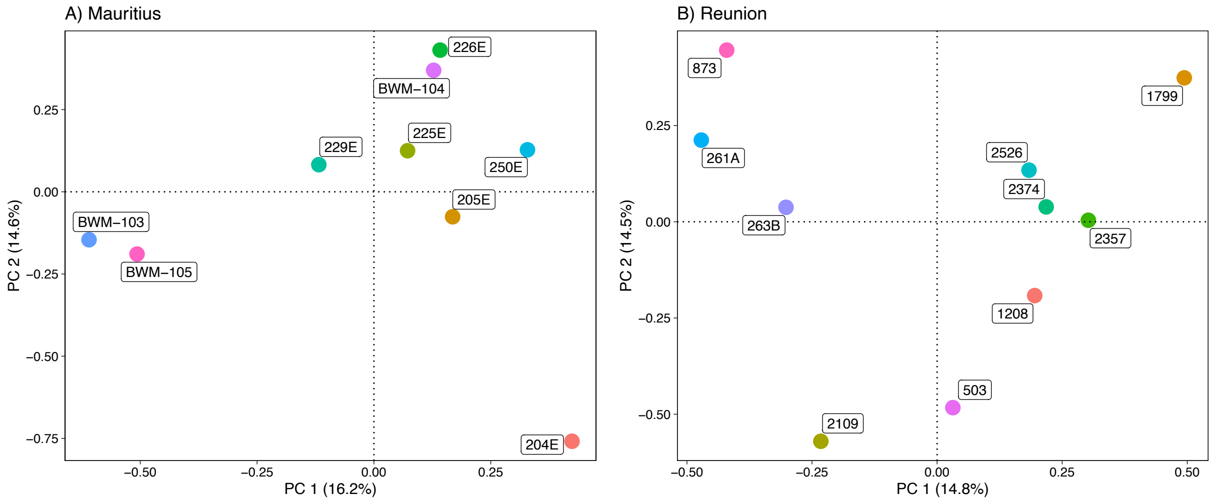

**Figure S11:** Likelihood distributions for the eight models tested in *fastsimcoal2* obtained from100 simulations of ML parameters. We exclude MOD1 for better visualization as its likelihood distribution is significantly lower (see Table S2). For MOD6_MIG_TR we present the two best runs that exemplify the contrasting relationships between current deme size and translocation rates ($H_NL_T$ and $L_NH_T$), see the Main text.

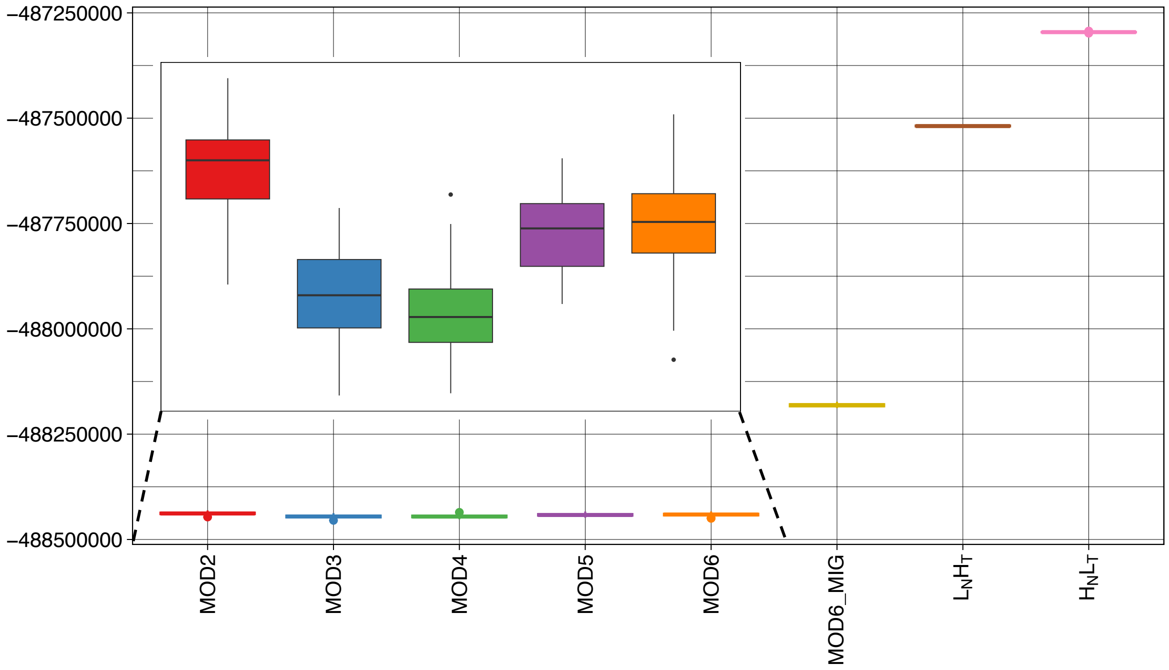

**Figure S12:** Scatter plot illustrating the relationship between $N_{MAU}$ and $TR_{MAD \rightarrow MAU}$ for eachof the 200 runs of MOD6_MIG_TR. Each point represents the outcome of a single run, colored according to the maximum estimated log-likelihood (MaxEstLhood). Histograms show the distribution of $N_{MAU}$ and $TR_{MAD \rightarrow MAU}$ across the 200 runs.

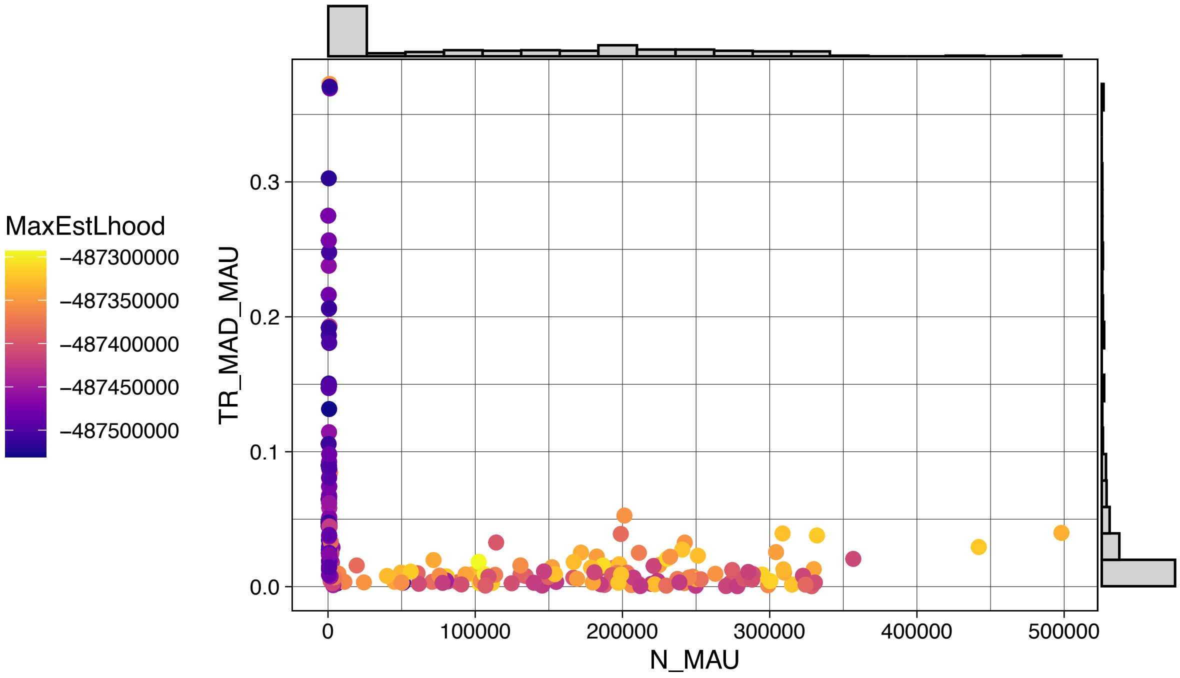

**Figure S13:** Scatter plot illustrating the relationship between $N_{REU}$ and $TR_{MAD→REU}$ for each ofthe 200 runs of MOD6_MIG_TR. Each point represents the outcome of a single run**,** coloredaccording to its maximum estimated log-likelihood (MaxEstLhood). Histograms show thedistribution of $N_{REU}$ and $TR_{MAD→REU}$ across the 200 runs.

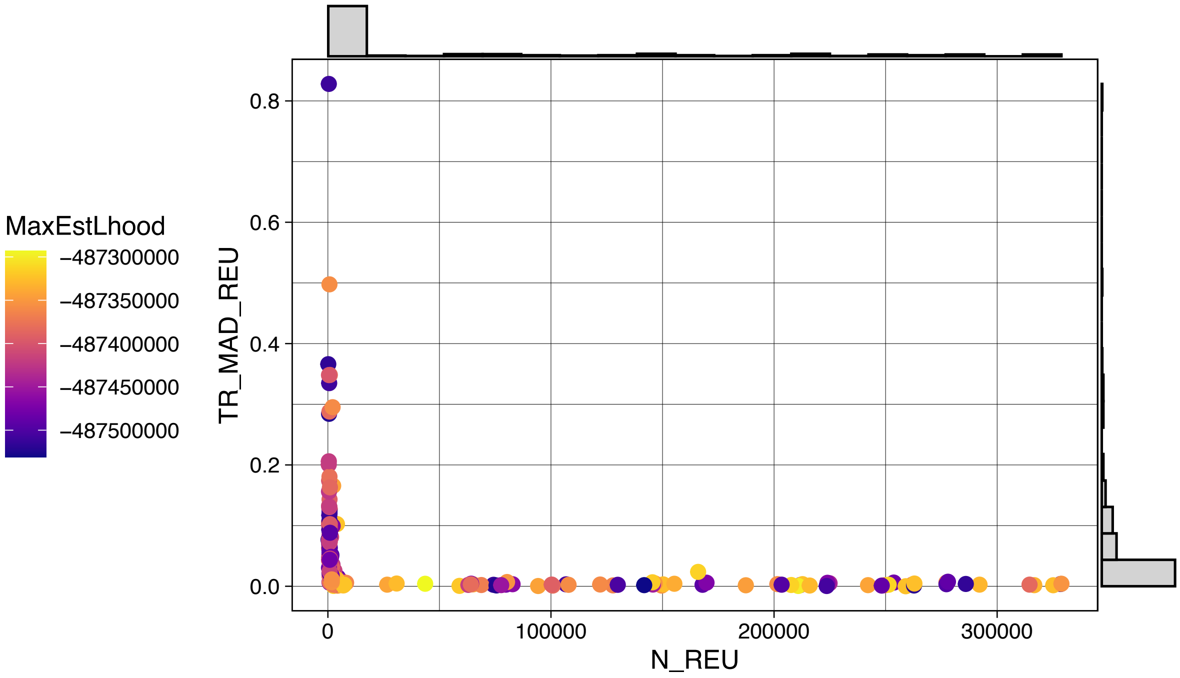

**Figure S14:** Density PCA plots generated from simulations of the parametric bootstraps for runs $H_NL_T$ and $L_NH_T$ of MOD6_MIG_TR. Each plot contains 100 points per population, where each point represents the barycenter of a population from a single simulation. The panels display results for: (A) HNLT and (B) LNHT. **(A)** $H_NL_T$. **(B)** $L_NH_T$.

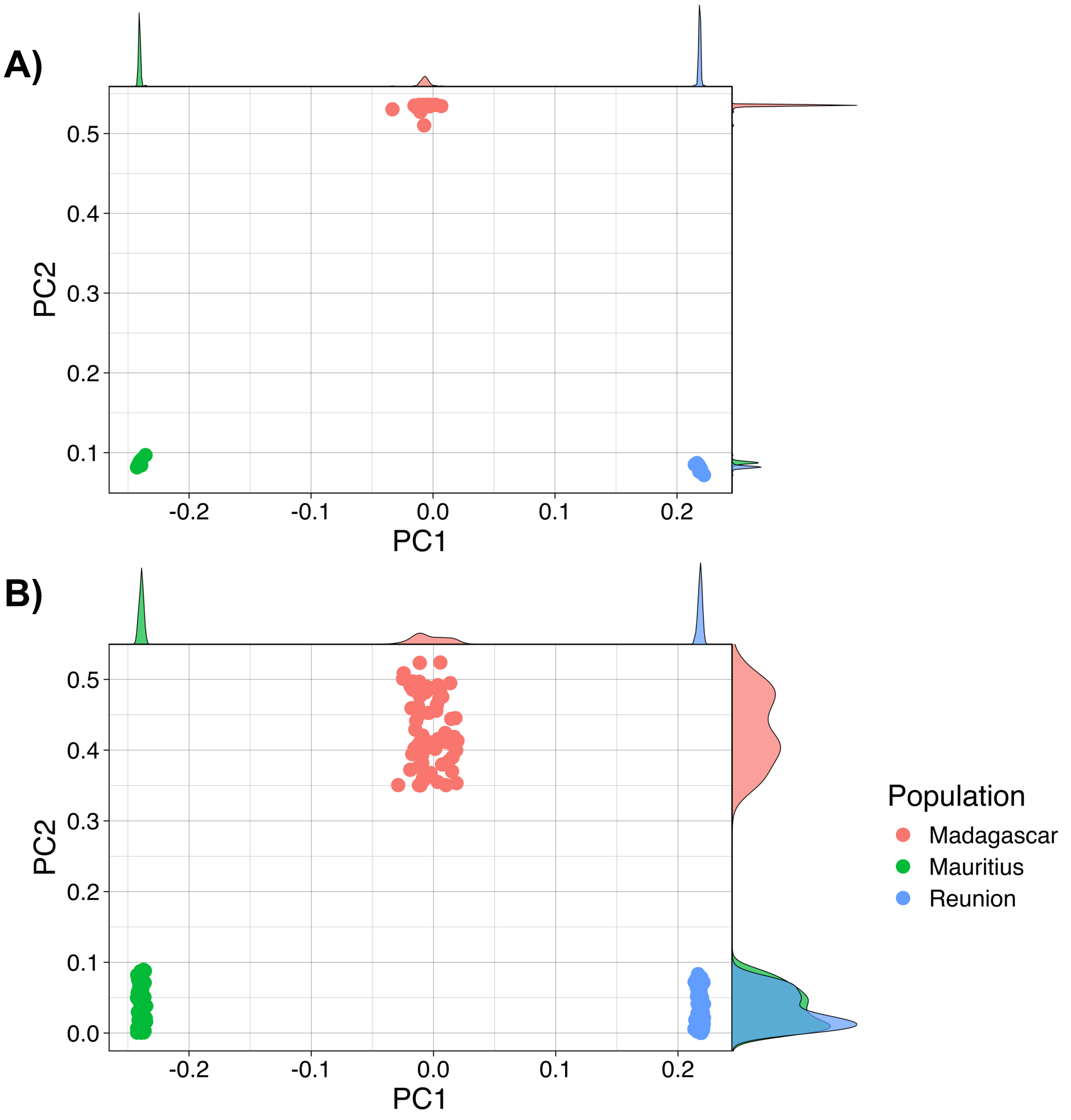

**Figure S15:** Pairwise Fst obtained from simulations of parametric bootstraps for $H_NL_T$ and $L_NH_T$ (MOD6_MIG_TR). Labels represent the mean pairwise Fst of each run and its 95% CI. The runs shown are: **(A)** $H_NL_T$. **(B)** $L_NH_T$.

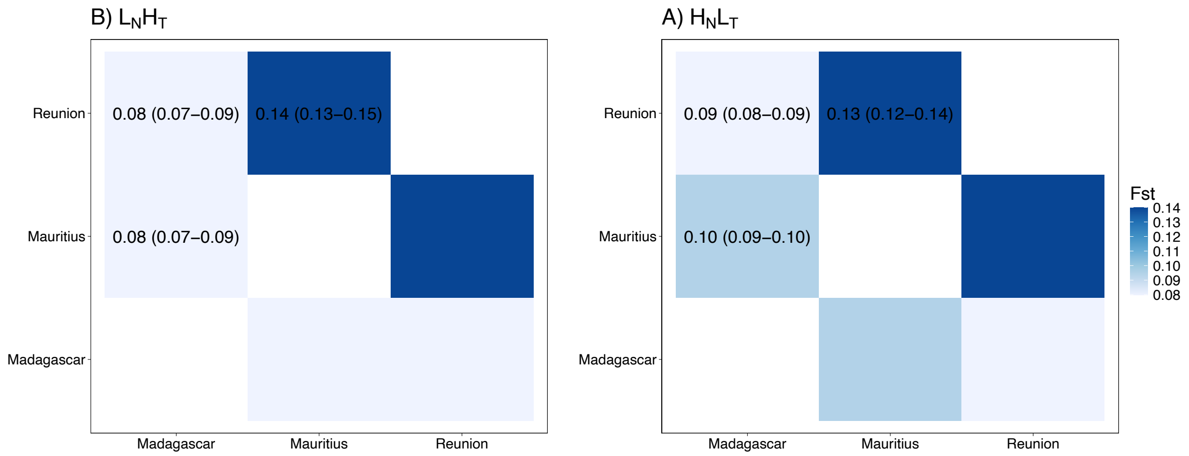

**Figure S16:** SNMF clustering of individuals simulated using the parametric bootstraps of $H_NL_T$ and $L_NH_T$ (MOD6_MIG_TR) according to the number of ancestral populations (K). The mean ancestry proportion was calculated from 100 simulations of each run. Clustering was done for: **(A)** Simulations of $H_NL_T$ with K=2. **(B)** Simulations of $H_NL_T$ with K=3. **(C)** Simulations of $L_NH_T$ with K=2. **(D)** Simulations of $L_NH_T$ with K=3.

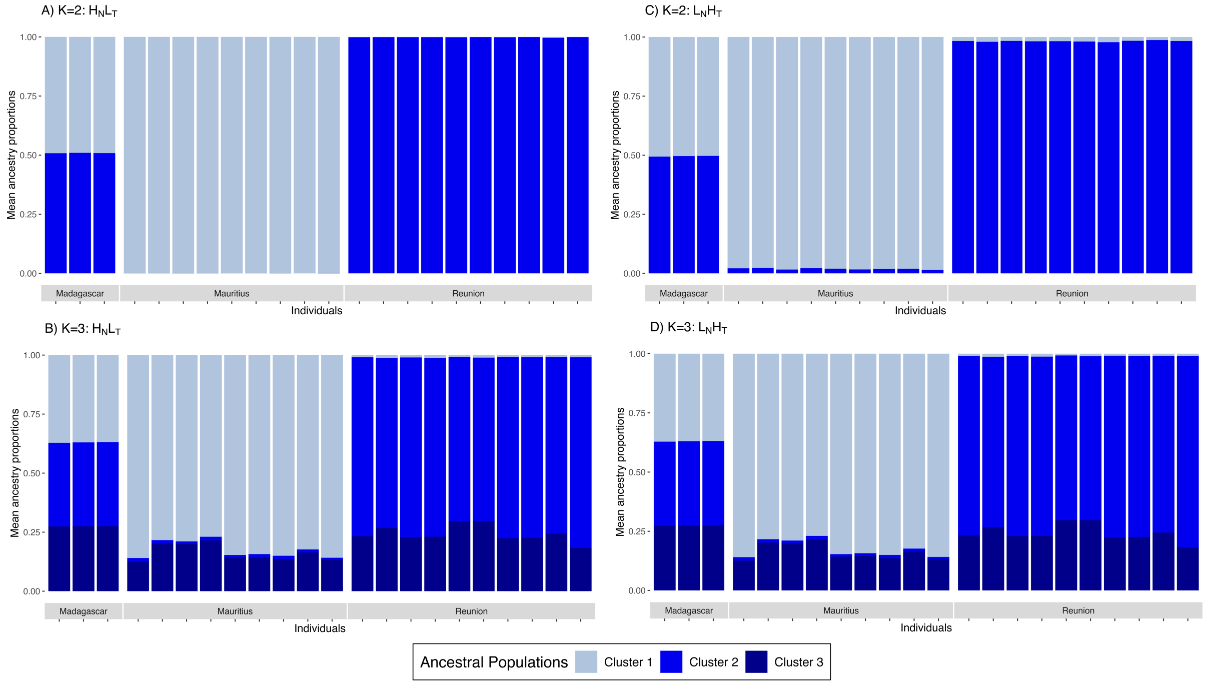

**Figure S17**: Comparison of genetic diversity statistics simulated from the parametric bootstraps of $H_N L_T$ and $L_N H_T$ (MOD6_MIG_TR). Error bars indicate the standard deviation around the mean across 100 simulations for each run. The statistics shown are: **(A)** Tajima's D. **(B)** $\theta_\pi$. **(C)** $\theta_W$.

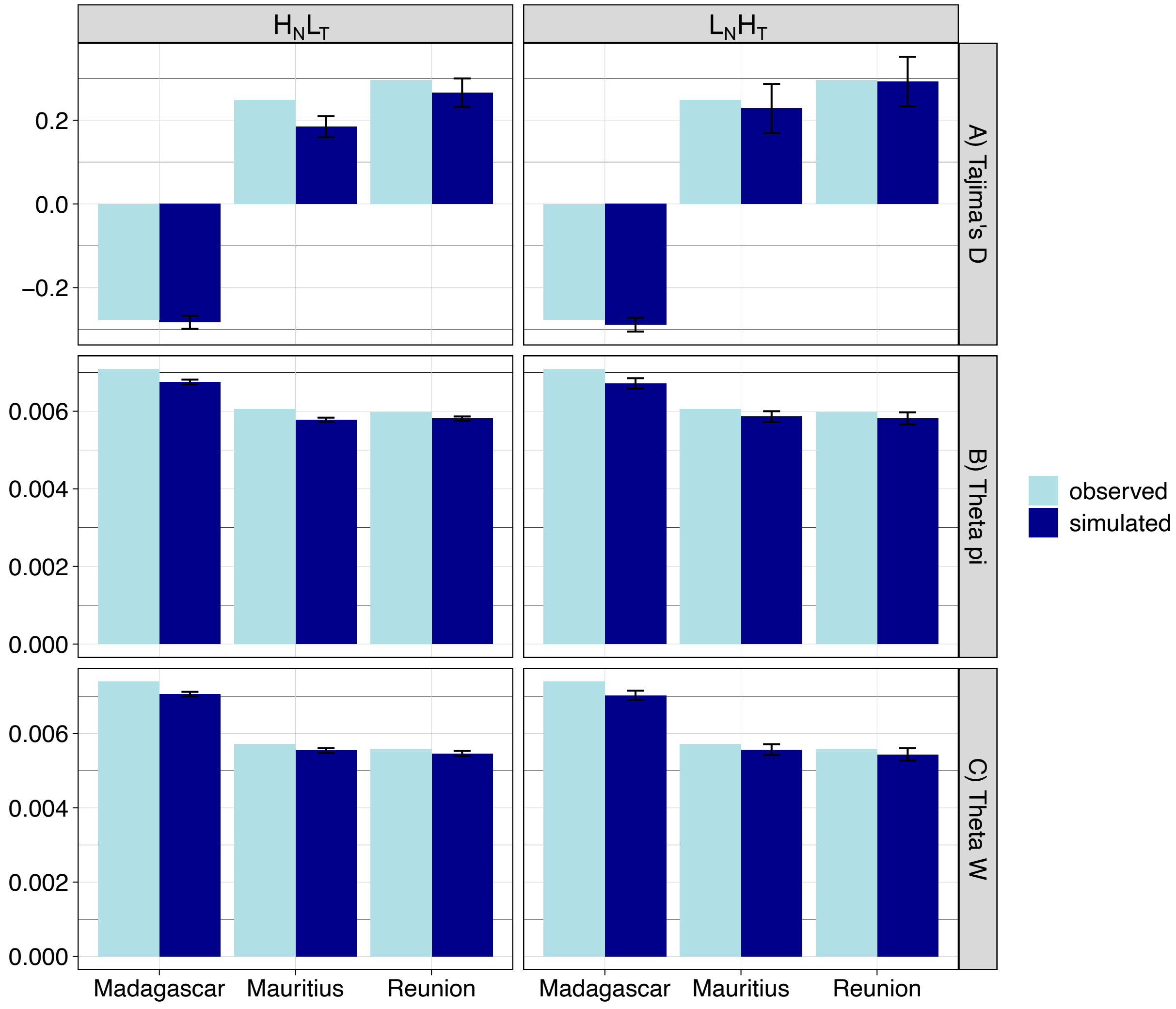

**Figure S18**: Boxplots showing the distribution of genetic diversity over 100k bp non-overlapping windows of scaffold1 for each population. The statistics shown are: **(A)** $\theta_{\pi}$. **(B)** $\theta_{W}$. **(C)** Tajima's D.

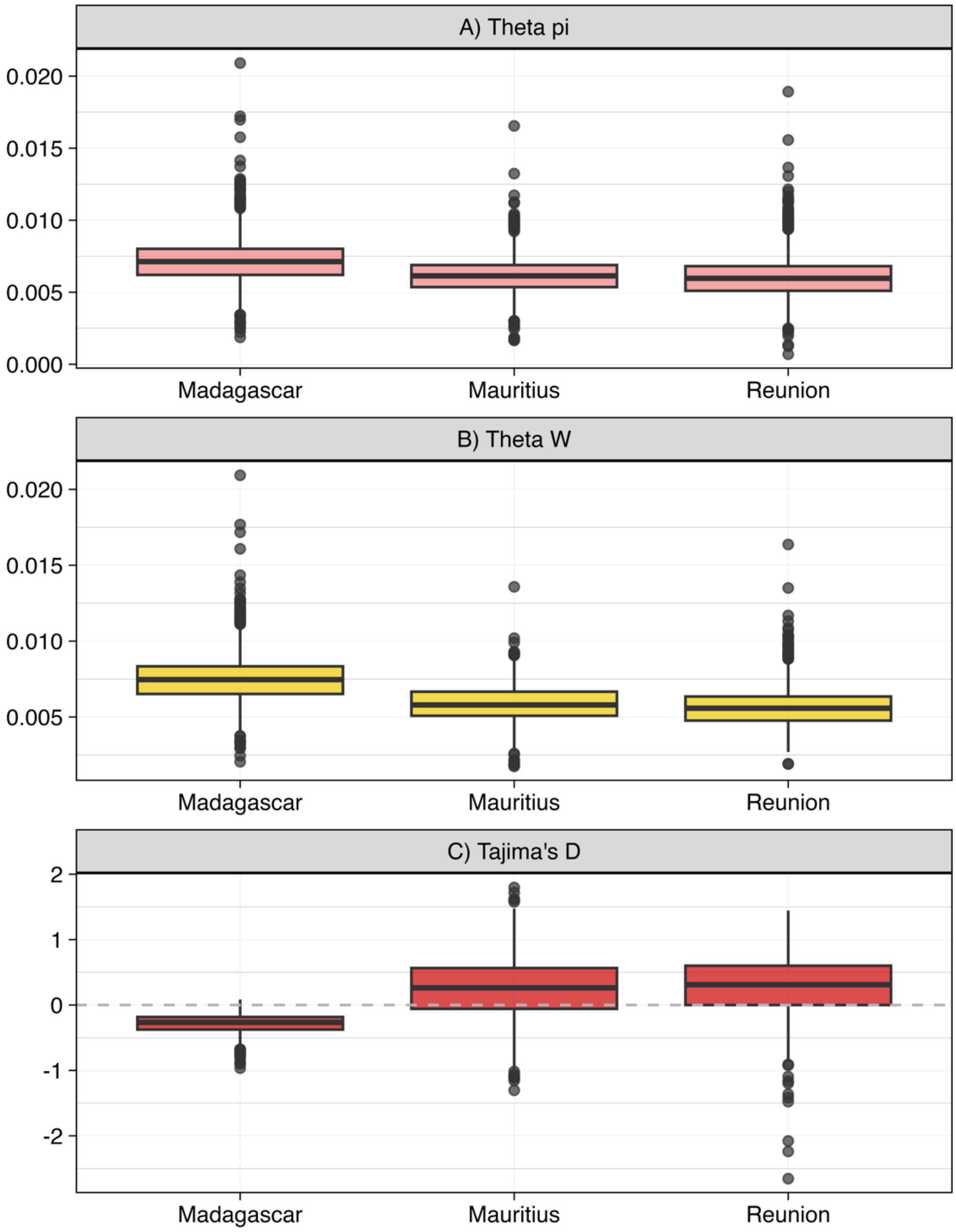

**Figure S19:** Simulated IICR from parametric bootstraps of $H_NL_T$ and $L_NH_T$ ofMOD6_MIG_TR.

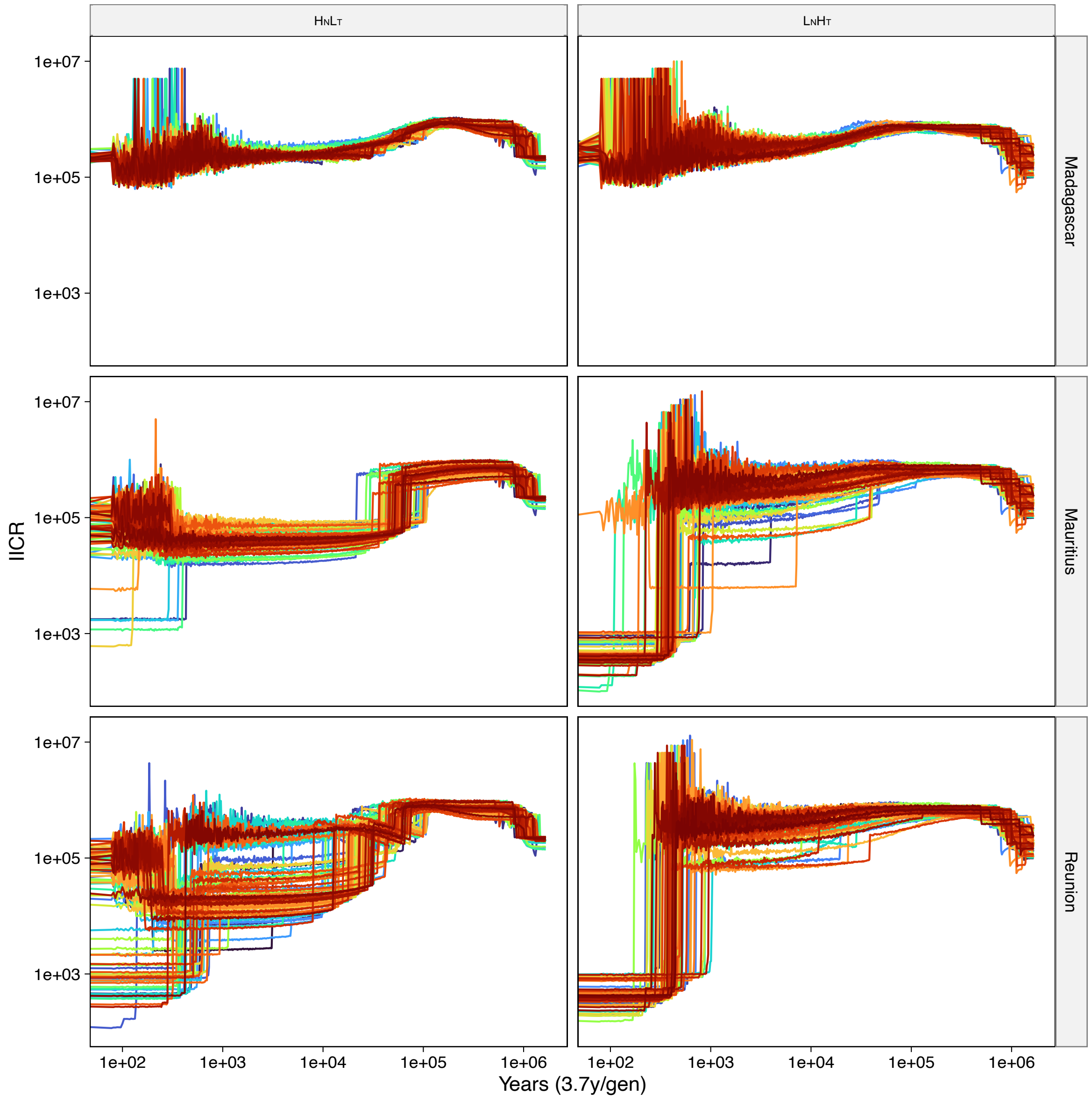

**Figure S20:** Comparing the IICR simulated from the ML parameters of $H_NL_T$ and $L_NH_T$ with the IICR inferred with PSMC and SMC++ from data simulated from both runs of MOD6_MIG_TR.

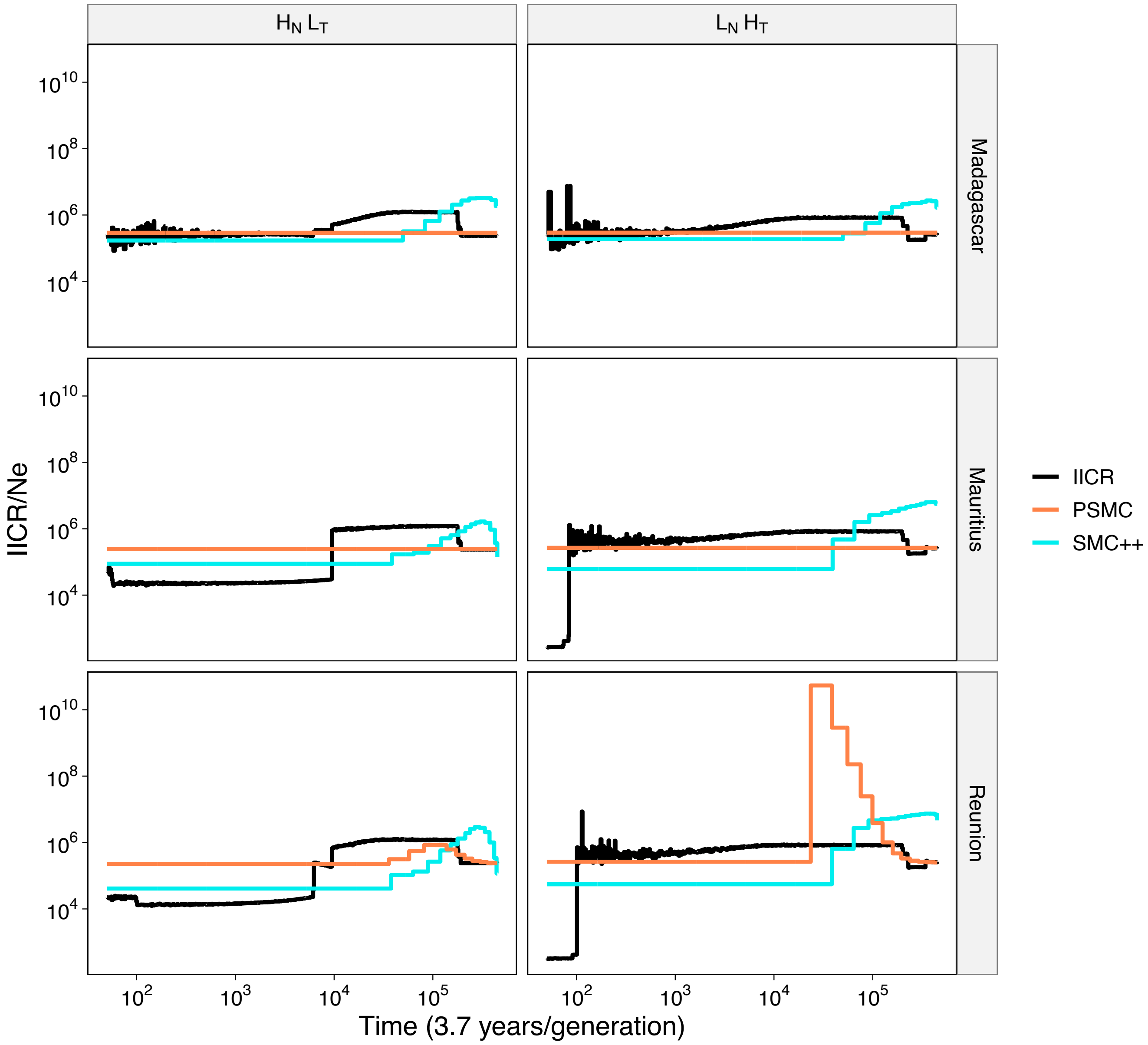

**Table S1:** Subfossils used for ancient DNA sequencing. All subfossils are from Mauritius. Sub1, Sub3 and Sub4 are from the collection of Julian P. Hume. Sub13 is from the MNHN paleontology collection. Species identifications were performed by JPH, based on bone morphology. calYBP, calender years before present. calAD, calendar years AD. ORAU, Oxford Radiocarbon Accelerator Unit.

| Lab specimen name | Family | Genus | Species | Site | Collector | Year of collection | Material | Age estimate | Source of age estimate | Percentage endogenous DNA | |
|---|---|---|---|---|---|---|---|---|---|---|---|
| | | | | | | | | | | Before capture | After capture |
| Sub1 | Columbidae | *Nesoenas* | Likely *cicur* or *picturata* | Mare aux Songes marsh, basin I | J. Hume | 2006 | Radius | 4200 calYBP | Radiocarbon dating of fossil layer, yielding dates consistently in range of 4100 - 4235 calYBP (Hume et al. 2015) | 0.285 | 1.501 |
| Sub4 | Columbidae | *Nesoenas* | *cicur* or *picturata* | Le Morne mountain | J. Hume | 2001 | Scapula | 320-134 calYBP (1705-1891 calAD) 95.4% probability | ORAU radiocarbon dating of subfossil, and calibration of date of 148 ± 19 radiocarbon YBP | 0.03 | 0.105 |
| Sub3 | Columbidae | *Nesoenas* | *cicur* or *picturata* | Le Pouce mountain, undercut in boulder scree | J. Hume | 2008 | Ulna | 200-1000 YBP* | Weathering of subfossil and identity of other fossils in fossil layer, representing endemic species and absence of exotic species | 1.516 | 4.19 |
| Sub13 | Columbidae | *Nesoenas* | Likely *picturata* | Le Pouce mountain | E. Thirioux | 1910 | Tarsometatarsus | 200-1000 YBP | Weathering of subfossil and time taken for subfossilisation prior to collection date | 3.053 | 5.834 |

*Radiocarbon dating of Sub3 was attempted by ORAU, but unfortunately it did not yield a date due to contamination of the subfossil with hardener used to protect it when returning from the field.

**Table S2:** Samples and metadata. All species attributions, including of subfossils, are based on morphology.

| Sample | Species | Population | Age group | Element | Mapped reads | Mean quality | Mean depth | Mean coverage (%) | Mean depth (scaffolds > 10 Mbp) | Mean coverage (%) (scaffolds > 10 Mbp) |
|---|---|---|---|---|---|---|---|---|---|---|
| 345630 | *N. picturata* | Madagascar | Modern | Blood | 293,915,495 | 36.1 | 35.9 | 77.0 | 32.2 | 99.5 |
| 436498 | *N. picturata* | Madagascar | Modern | Tissue | 514,527,544 | 36.3 | 33.9 | 77.7 | 38.4 | 99.7 |
| 438653 | *N. picturata* | Madagascar | Modern | Tissue | 367,237,931 | 36.1 | 17.8 | 75.6 | 28.0 | 99.5 |
| BWM-103 | *N. picturata* | Mauritius | Modern | Blood | 265,924,516 | 35.9 | 19.3 | 75.1 | 29.4 | 99.6 |
| BWM-104 | *N. picturata* | Mauritius | Modern | Blood | 253,784,758 | 36 | 15.3 | 73.2 | 28.7 | 99.5 |
| BWM-105 | *N. picturata* | Mauritius | Modern | Blood | 354,925,451 | 36.1 | 29.9 | 78.3 | 39.4 | 99.7 |
| 204E | *N. picturata* | Mauritius | Modern | Blood | 462,152,637 | 36.3 | 20.3 | 79.0 | 35.0 | 99.7 |
| 205E | *N. picturata* | Mauritius | Modern | Blood | 639,821,009 | 36.1 | 23.6 | 79.8 | 49.0 | 99.7 |
| 225E | *N. picturata* | Mauritius | Modern | Blood | 590,549,916 | 36.5 | 20.8 | 77.7 | 45.8 | 99.8 |
| 226E | *N. picturata* | Mauritius | Modern | Blood | 366,929,487 | 36.3 | 13.0 | 75.4 | 28.7 | 99.5 |
| 229E | *N. picturata* | Mauritius | Modern | Blood | 518,882,468 | 36.3 | 23.4 | 78.7 | 39.5 | 99.8 |
| 250E | *N. picturata* | Mauritius | Modern | Blood | 506,464,365 | 36.3 | 15.7 | 75.7 | 39.7 | 99.7 |
| 1208 | *N. picturata* | Reunion | Modern | Tissue | 432,826,252 | 36.3 | 27.5 | 79.4 | 33.0 | 99.7 |
| 1799 | *N. picturata* | Reunion | Modern | Tissue | 465,980,084 | 36.3 | 25.0 | 79.2 | 35.4 | 99.7 |
| 2109 | *N. picturata* | Reunion | Modern | Tissue | 500,579,847 | 36.3 | 28.3 | 78.8 | 38.1 | 99.7 |
| 2357 | *N. picturata* | Reunion | Modern | Tissue | 475,158,583 | 36.3 | 33.5 | 78.9 | 36.5 | 99.6 |
| 2374 | *N. picturata* | Reunion | Modern | Tissue | 416,809,638 | 36.3 | 20.7 | 79.1 | 31.8 | 99.6 |
| 2526 | *N. picturata* | Reunion | Modern | Tissue | 520,015,809 | 36.4 | 27.2 | 79.2 | 40.3 | 99.7 |
| 503 | *N. picturata* | Reunion | Modern | Tissue | 470,020,628 | 36.3 | 29.6 | 79.6 | 36.6 | 99.7 |
| 873 | *N. picturata* | Reunion | Modern | Tissue | 263,746,651 | 36.1 | 23.3 | 75.2 | 28.8 | 99.4 |
| 261A | *N. picturata* | Reunion | Modern | Blood | 485,415,850 | 36.2 | 15.5 | 74.5 | 37.1 | 99.7 |
| 263B | *N. picturata* | Reunion | Modern | Blood | 411,799,732 | 36.3 | 14.1 | 74.4 | 32.2 | 99.7 |
| Sub1 | *N. cicur/ picturata* | Mauritius | Subfossil | Radius | 134,806 | 38.2 | 0.004 | 0.3 | | |
| Sub3 | *N. cicur/ picturata* | Mauritius | Subfossil | Ulna | 2,719,895 | 38.5 | 0.041 | 2.8 | | |
| Sub4 | *N. cicur/ picturata* | Mauritius | Subfossil | Scapula | 31,024 | 40.5 | 0.002 | 0.1 | | |
| Sub13 | *N. picturata* | Mauritius | Subfossil | Tarsometatarsus | 1,821,912 | 40.4 | 0.031 | 2.8 | | |

**Table S3**: ML parameter estimates for the eight models tested in *fastsimcoal2* with associated distributions and ranges. Times are expressed in years and migration rates are indicated forward in time.

| Parameter | Parameter distribution | Parameter range | MOD1 | MOD2 | MOD3 | MOD4 | MOD5 | MOD6 | MOD6_MIG | MOD6_MIG_TR_RUN92 | MOD6_MIG_TR_RUN113 |
|---|---|---|---|---|---|---|---|---|---|---|---|
| $N_{MAU}$ | uniform | 100-400,000 | 372,724 | 206,911 | 209,767 | 219,904 | 204,937 | 234,172 | 34,527 | 102,403 | 540 |
| $N_{REU}$ | uniform | 100-400,000 | 262,632 | 184,024 | 194,671 | 208,713 | 196,073 | 174,582 | 140,696 | 43,602 | 629 |
| $N_{MAD}$ | uniform | 100-400,000 | 183,173 | 637,737 | 634,759 | 630,541 | 636,153 | 629,106 | 1,122,452 | 479,042 | 512,869 |
| $N_{MAU1}$ | uniform | 100-400,000 | | | | | | | | 43,227 | 532,314 |
| $N_{REU1}$ | uniform | 100-400,000 | | | | | | | | 26,150 | 617,752 |
| $N_{MAU2}$ | uniform | 100-400,000 | | | | | | | | 1,394,066 | 53,309 |
| $N_{REU2}$ | uniform | 100-400,000 | | | | | | | | 608,445 | 178,437 |
| $N_{ANC}$ | uniform | 100-400,000 | 463,130 | | | | | | | | |
| T1 | uniform | 100-200,000 | 269,860 | 119,036 | 120,972 | 126,414 | 120,143 | 136,145 | 793,376 | 659,514 | 1,261,478 |
| T2 | uniform | 100-200,000 | 229,370 | 110,612 | 119,884 | 124,139 | 118,541 | 101,025 | 2,024,588 | 698,623 | 1,449,494 |
| T3 | uniform | 100-200,000 | | | | | | | | 34,787 | 836,167 |
| T4 | uniform | 100-200,000 | | | | | | | | 22,718 | 736,996 |
| T5 | uniform | 100-200,000 | | | | | | | | 207 | 307 |
| T6 | uniform | 100-200,000 | | | | | | | | 366 | 370 |
| T7 | uniform | 1-100 | | | | | | | | 170 | 270 |
| T8 | uniform | 1-100 | | | | | | | | 329 | 333 |
| $m_{REU \rightarrow MAU}$ | uniform | 0-0.0001 | | | | | | | 1.01E-06 | 1.88E-06 | 7.42E-05 |
| $m_{MAD \rightarrow MAU}$ | uniform | 0-0.0001 | | | | | | | 6.61E-05 | 1.33E-05 | 1.56E-05 |
| $m_{MAU \rightarrow REU}$ | uniform | 0-0.0001 | | | | | | | 1.37E-05 | 4.17E-05 | 6.67E-05 |
| $m_{MAD \rightarrow REU}$ | uniform | 0-0.0001 | | | | | | | 1.84E-07 | 4.70E-06 | 7.79E-05 |
| $m_{MAU \rightarrow MAD}$ | uniform | 0-0.0001 | | | | | | | 1.91E-06 | 1.44E-05 | 1.96E-05 |
| $m_{REU \rightarrow MAD}$ | uniform | 0-0.0001 | | | | | | | 8.85E-06 | 3.03E-05 | 8.28E-05 |
| $TR_{MAD \rightarrow MAU}$ | uniform | 0-0.9 | | | | | | | | 0.018393 | 0.1862601 |
| $TR_{MAD \rightarrow REU}$ | uniform | 0-0.9 | | | | | | | | 0.0041394 | 0.1170953 |
| MaxEstLhood | | | -954,932,002 | -488,405,652 | -488,418,645 | -488,412,981 | -488,411,331 | -488,415,308 | -487,603,871 | -487,293,304 | -487,510,379 |
| MaxObsLhood | | | -487,164,528 | -487,164,528 | -487,164,528 | -487,164,528 | -487,164,528 | -487,164,528 | -487,164,528 | -487,164,528 | -487,164,528 |
| AIC | | | 4,397,624,396 | 2,249,191,155 | 2,249,250,994 | 2,249,224,909 | 2,249,217,309 | 2,249,235,625 | 2,245,498,833 | 2,244,068,638 | 2,245,068,304 |
